# Subgenome dominance shapes novel gene evolution in the decaploid pitcher plant *Nepenthes gracilis*

**DOI:** 10.1101/2023.06.14.544965

**Authors:** Franziska Saul, Mathias Scharmann, Takanori Wakatake, Sitaram Rajaraman, André Marques, Matthias Freund, Gerhard Bringmann, Louisa Channon, Dirk Becker, Emily Carroll, Yee Wen Low, Charlotte Lindqvist, Kadeem J. Gilbert, Tanya Renner, Sachiko Masuda, Michaela Richter, Gerd Vogg, Ken Shirasu, Todd P. Michael, Rainer Hedrich, Victor A. Albert, Kenji Fukushima

## Abstract

Subgenome dominance after whole-genome duplication generates distinction in gene number and expression at the level of chromosome sets, but it remains unclear how this process may be involved in evolutionary novelty. Here, we generated a chromosome-scale genome assembly of the Asian pitcher plant *Nepenthes gracilis* to analyze how its novel traits (dioecy and carnivorous pitcher leaves) are linked to genomic evolution. We found a decaploid karyotype with five complete sets of syntenic chromosomes (2*n* = 10*x* = 80) yet with a clear indication of subgenome dominance and highly diploidized gene contents. The male-linked and pericentromerically located region on the putative sex chromosome was identified in a recessive subgenome and was found to harbor three transcription factors involved in flower and pollen development, including a likely neofunctionalized *LEAFY* duplicate. Transcriptomic and syntenic analyses of carnivory-related genes suggested that the paleopolyploidization events seeded genes that subsequently formed tandem clusters in recessive subgenomes with specific expression in the digestive zone of the pitcher, where specialized cells digest prey and absorb derived nutrients. Novel gene evolution in recessive subgenomes is likely to be prevalent because duplicates were enriched with *Nepenthes*-specific genes with tissue-specific expression, including those expressed in trapping pitchers. Thus, subgenome dominance likely contributed to evolutionary novelty by allowing recessive subgenomes experiencing relaxed purifying selection to serve as a preferred host of novel tissue-specific duplicates. Our results provide insight into how polyploids, which may frequently be evolutionary dead-ends, have given rise to novel traits in exceptionally thriving high-ploidy lineages.

## Introduction

Novel phenotypes are often the result of the emergence of new genes that have undergone functional divergence after gene duplication. Polyploidy, which involves the duplication of entire genomes, has been argued to have great potential to drive adaptive evolution (Soltis et al., 2009). However, polyploid lineages are sometimes considered evolutionary dead-ends, with only short-term adaptive potential and high long-term extinction rates (Van de Peer et al., 2017). There are, however, some exceptions to this trend, such as flowering plants (angiosperms) (AMBORELLA GENOME PROJECT et al., 2013; Chanderbali et al., 2022) and vertebrates (Dehal and Boore, 2005), which have thrived following multiple rounds of polyploidization early in their evolution.

After polyploidization, especially following a hybridization event (Edger et al., 2017; Li et al., 2021), the coexistence of more than one haploid chromosome set often engenders subgenome dominance, in which one chromosome set is preferentially expressed over the other (Cheng et al., 2018). This is one of the early changes that occur after polyploidization, and the differential retention of duplicate genes on dominant versus recessive subgenomes can render versatile adaptive prospects between these subgenomes over time. Although such a genome-wide effect may constrain the long-term evolutionary paths of a large number of genes, it remains unclear how subgenome dominance may be involved in the evolution of novel gene functions. To study the long-term effects of subgenome dominance, it is necessary to examine genomes including relatively ancient polyploidization events, yet with well-preserved subgenome structures.

The Old World pitcher plant *Nepenthes* offers a unique opportunity to study the relationship between polyploidization-related genomic features and the evolution of novel genes. This genus of carnivorous plants includes at least 169 species of perennial vines (Cross et al., 2020) and is known for its highly modified leaves (pitchers) adapted for trapping and obtaining nutrients from arthropod prey (Ellison and Adamec, 2018). In addition to carnivory, *Nepenthes* possess other atypical traits, such as dioecy. Less than 1% of all angiosperms are carnivorous (Cross et al., 2020), and approximately 6% of angiosperms are dioecious, possessing distinct male and female individuals (Renner and Ricklefs, 1995). No carnivorous plant species outside of *Nepenthes* are known to be dioecious. Thus, no other plant lineages possess this unique combination of traits. *Nepenthes* has undergone ancient whole-genome duplications (WGDs), as indicated by transcriptomic signatures (Walker et al., 2017; Yang et al., 2018; Palfalvi et al., 2020), and exhibits highly stable karyotypes among many species (Heubl and Wistuba, 1997).

In this study, we present a chromosome-scale genome assembly of *Nepenthes gracilis* and demonstrate that the *Nepenthes* genome is decaploid, bearing five subgenomes with a clear signature of subgenome dominance. Our analysis suggests that recessive subgenomes played a crucial role in permitting the adaptive evolution of tissue-specific genes associated with the unique biology of *Nepenthes*, and we discuss how gene divergence occurs under the influence of WGDs, subgenome dominance, and subsequent small-scale duplications (SSDs).

## Results

### Genome assembly and annotation

We used a combination of Oxford Nanopore Technology (ONT) long-read sequencing and Illumina short-read sequencing (Supplementary Table 1) to generate megabase-scale genome assemblies for male and female *Nepenthes gracilis* plants (Fig. 1a; Supplementary Table 2), whose *k*-mer-based genome size estimates were around 722 Mb (Supplementary Fig. 1). The male assembly was further improved through Hi-C scaffolding to generate a 752.9-Mb genome assembly containing 40 chromosome-scale scaffolds (N50 = 18.6 Mb), which accounted for 99.2% (746.7 Mb) of the total assembly size (Supplementary Fig. 2), consistent with the reported number of chromosomes in this species (Heubl and Wistuba, 1997). Unless otherwise noted, we report the analysis of the male Hi-C assembly in this work. Repetitive elements accounted for 67.17% of the reference genome (Supplementary Table 3). Using RNA-seq data (Supplementary Table 4), we predicted a total of 34,010 gene models (Supplementary Table 5) with a gene-space completeness score of 94.1% (1,519/1,614 genes in the embryophyta_odb10 dataset) as determined by Benchmarking Universal Single-Copy Orthologs (BUSCO) (Manni et al., 2021) (Supplementary Table 6).

**Figure 1.**
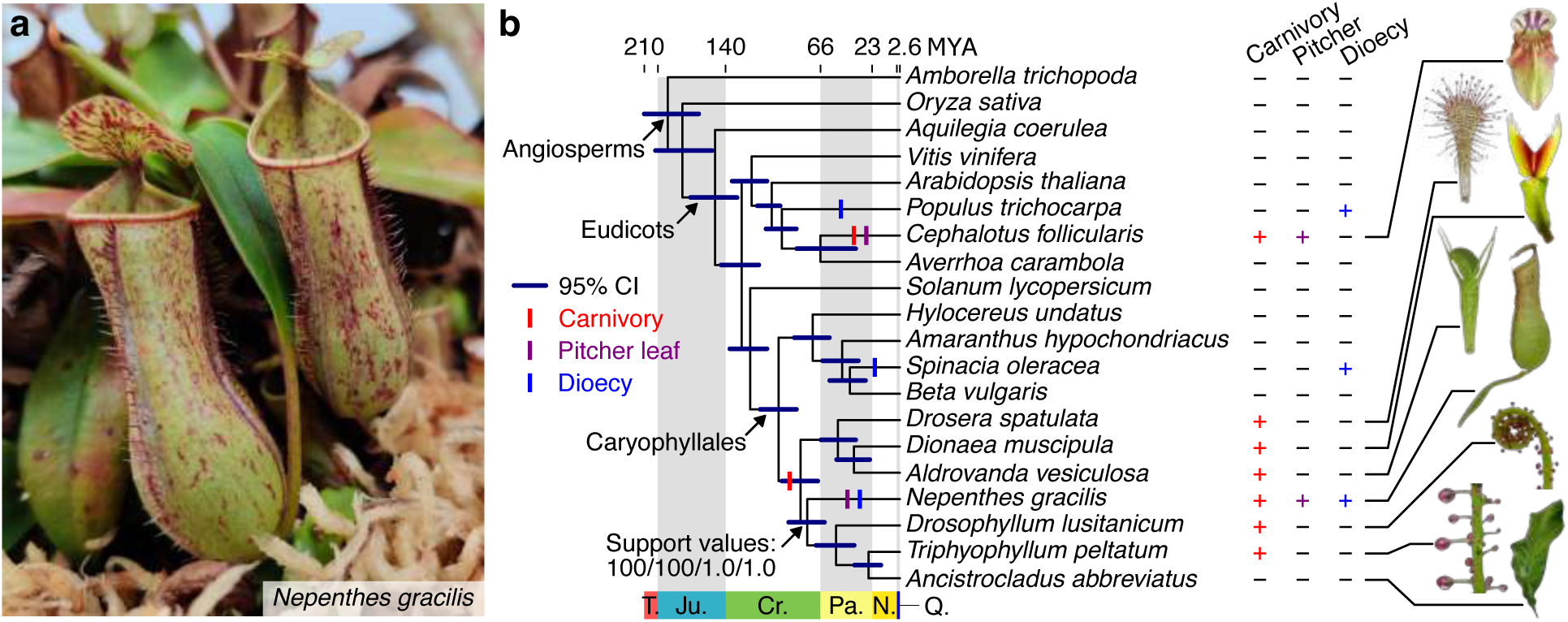
Evolution of novel traits in *Nepenthes*. (**a**) *Nepenthes gracilis* with carnivorous pitcher leaves. (**b**) The phylogenetic position of *Nepenthes*. Divergence time with a 95% confidence interval (CI) is shown for the maximum-likelihood (ML) tree topology reconstructed using 1,614 single-copy protein sequences. See Supplementary Fig. 3 for results with alternative phylogenetic methods, including the coalescence-based (CO) approach. Bootstrap supports and posterior probability values for the position of *Nepenthes* are shown as follows: nucleotide ML / protein ML / nucleotide CO / protein CO. Character evolution was parsimoniously mapped to branches, and symbols do not indicate the point estimate of evolutionary origins. Note that carnivory was secondarily lost in *Ancistrocladus* (Heubl et al., 2006). Leaves of plants belonging to carnivorous clades are shown to the right. CI, confidence interval; MYA, million years ago; T., Triassic; Ju., Jurassic; Cr., Cretaceous; Pa., Paleogene; N., Neogene; Q., Quaternary.

### Character evolution under a robust phylogeny

The phylogenetic position of *Nepenthes* and its monotypic family, Nepenthaceae, has been a matter of debate, even with the use of improved gene-tree mining from transcriptomic data. Previous phylogenetic hypotheses have placed Nepenthaceae as a sister to Droseraceae, to a clade of three families (Ancistrocladaceae, Dioncophyllaceae, and Drosophyllaceae; ADD families), or to a clade containing all four of these plant families (Supplementary Fig. 3a–c). To more accurately determine the phylogenetic position of Nepenthaceae, we performed phylogenomic analyses using gene models from the newly sequenced *Nepenthes* genome. We also sequenced the transcriptomes of the ADD families, for which genome sequences are currently unavailable (Supplementary Table 6). Our analysis, using 1,614 Embryophyta-wide single-copy BUSCO genes (Supplementary Fig. 3d), resulted in the same tree topology, regardless of the inference method (maximum-likelihood (Minh et al., 2020) or coalescence-based (Zhang et al., 2018)) or the substitution model used (GTR+R4 for nucleotides or LG+R4 for amino acids) (Supplementary Fig. 3e). This tree topology, limited of course to a bifurcating evolutionary history (see Supplementary Text 1 for the possibility of ancient hybridization events), indicated that carnivorous plants in this lineage likely split in the sequential order of Droseraceae, Nepenthaceae, and then the rest. Under parsimony, these phylogenetic relationships support the hypothesis that carnivory evolved only once in Caryophyllales and that the pitfall-type trap leaves of Nepenthaceae were derived from flypaper-type trap leaves (Albert et al., 1992; Heubl et al., 2006; Freund et al., 2022), which are found in Droseraceae, Drosophyllaceae, and Dioncophyllaceae (Fig. 1b). Dioecy is another fascinating character evolved uniquely in Nepenthaceae after its split from the other lineages (Scharmann et al., 2019).

### Decaploidal origin of the *Nepenthes* genome

The large chromosome number in *Nepenthes* (*n* = 40) (Heubl and Wistuba, 1997) could suggest a history of WGDs. Although ancient WGDs have been reported in *Nepenthes* and related lineages based on synonymous substitution plots of duplicate gene pairs (Walker et al., 2017; Yang et al., 2018; Palfalvi et al., 2020), the exact ploidy level and its impact on *Nepenthes* genome structure have not yet been determined. To investigate polyploidization history in *Nepenthes*, we analyzed patterns of internal synteny, revealing that the haploid-level *Nepenthes* genome has a clear structure of eight syntenic groups, each containing exactly five chromosomes (Fig. 2a), indicating a decaploidal (five subgenomes) origin with a basic chromosome number of eight (i.e., *x* = 8). The distribution of synonymous distances between paralog pairs (Supplementary Fig. 4), coupled with clear 4:1 patterns of fractionation bias among homeologous chromosomes (Supplementary Fig. 5), indicates that *Nepenthes* has a complex polyploid background, having undergone at least two sequential WGDs following the *gamma* hexaploidization event that occurred in the common ancestor of all extant core eudicots (Jaillon et al., 2007). This history has also included the addition of a fifth subgenome at some point during the evolution of the *Nepenthes* lineage, possibly after splitting from its closest relatives (Supplementary Fig. 4). Given that the chromosome number is stable in *Nepenthes*, including in the earliest diverged species *N. pervillei* (Heubl and Wistuba, 1997), decaploidy must have been established before the diversification of extant species (6.4–18.2 MYA (Scharmann et al., 2021)). This suggests the possibility that these ancient WGDs may have played a role in the evolution of novel traits in *Nepenthes*.

**Figure 2.**
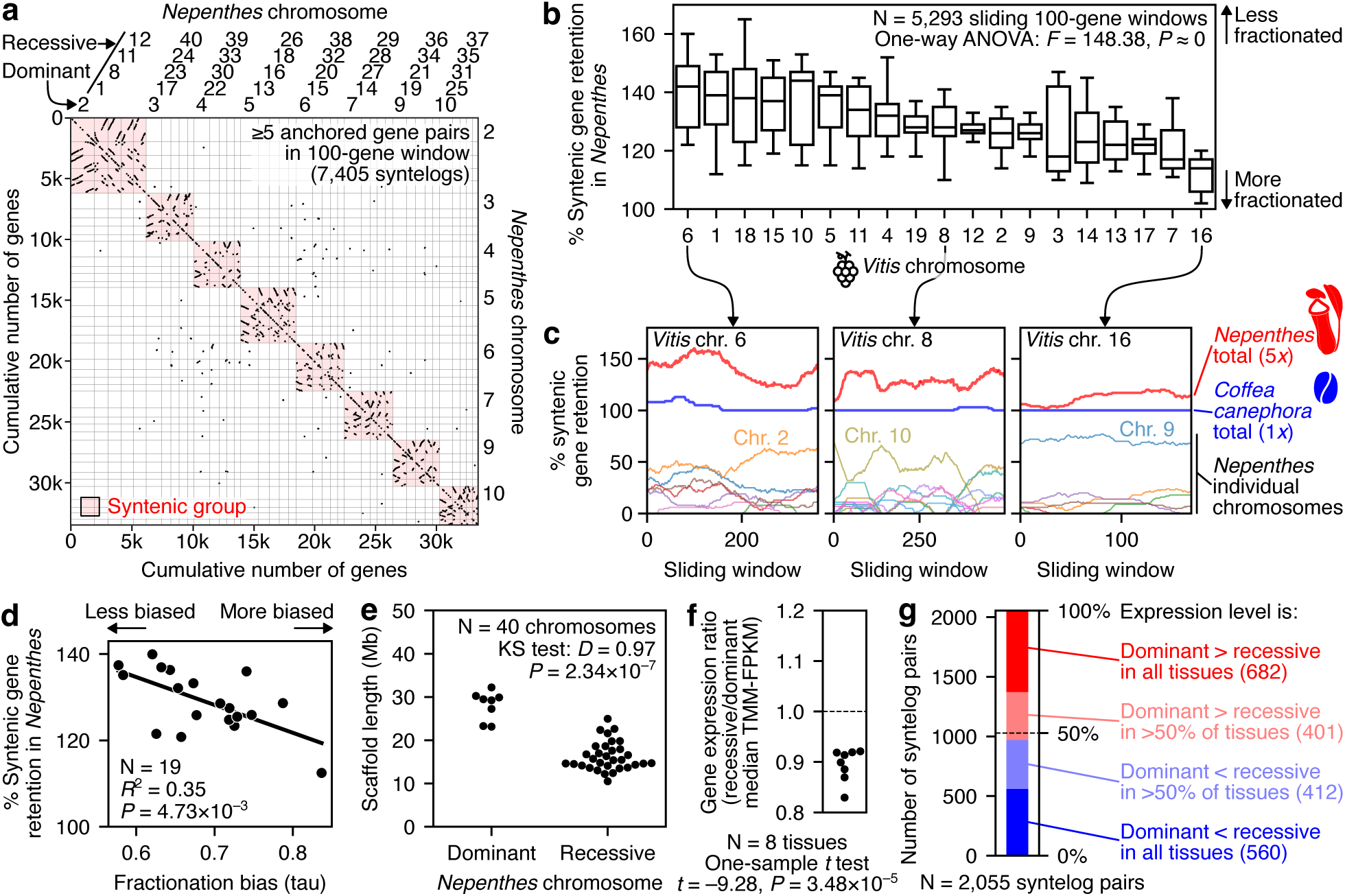
Subgenome dominance in the decaploidal *Nepenthes* genome. (**a**) Self-self syntenic dotplot of protein-coding sequences shows clear evidence for syntenic groups. Chromosomes are ordered by identified homeologous groups (red). For the identification of dominant/recessive chromosomes, see **c** and Supplementary Fig. 5. (**b**) The unequal rate of fractionation in the *Nepenthes* genome. Distribution of the syntenic gene retention rate is shown for syntenic blocks grouped by corresponding *Vitis* chromosomes. The statistical significance of unequal rates was tested with a one-way analysis of variance (ANOVA). Box plot elements are defined as follows: center line, median; box limits, upper and lower quartiles; whiskers, 1.5 × interquartile range. (**c**) Different levels of fractionation in the *Nepenthes* syntenic blocks (5*x*) mapped to representative *Vitis* chromosomes (1*x*): lowest in chromosome 6, middle in chromosome 8, and highest in chromosome 16. Analysis of the *Coffea canephora* genome (also 1*x*) with the same FractBias parameters (Joyce et al., 2017) is shown for comparison. (**d**) A negative correlation between fractionation bias and syntenic gene retention. Yanai’s tau (Yanai et al., 2005) was utilized as a proxy for fractionation bias. Linear regression and associated statistics are provided in the plot. (**e**) Dominant chromosomes tend to be larger than recessive ones. Statistical significance was examined with a Kolmogorov-Smirnov test. (**f**) Syntelogs in the dominant chromosomes tend to express at higher levels than those in the recessive chromosomes. Statistical significance was tested with a one-sample *t*-test. Eight tissues in *Nepenthes* were analyzed (Supplementary Table 4). TMM-FPKM: fragments per kilobase million normalized by the trimmed mean of M-values. (**g**) Syntelog-wise comparison of expression levels in genes on a dominant chromosome versus on a recessive chromosome.

### Biased gene fractionation and subgenome dominance

Despite having a complete decaploid karyotype (*n* = 5*x* = 40 and 2*n* = 10*x* = 80), the gene content in *Nepenthes* is highly fractionated, as is evident in a syntenic comparison with grape (*Vitis vinifera*), which has maintained its ploidy level since the *gamma* hexaploidization event (i.e., 1*x* in *Vitis* versus 5*x* in *Nepenthes*) (Jaillon et al., 2007) (Fig. 2b). In total, 94.5% (5,002/5,293) of 100-gene genomic windows in the *Nepenthes* genome retained fewer than 1.5 syntelog copies on average (compared to the ploidy-based expectation of 5.0 copies). While fractionation is advanced overall, there is significant heterogeneity among syntenic chromosome groups (Fig. 2b). For example, the syntenic group corresponding to grape chromosome 6 has an average of 1.4 syntelog copies, while the group corresponding to grape chromosome 16 has an average of only 1.1 copies. Homeologous *Nepenthes* chromosome groups that are highly fractionated tend to show a clear distinction between one dominant, gene-rich chromosome and four recessive, gene-poor chromosomes (i.e., fractionation bias (Joyce et al., 2017)). This distinction is less obvious among groups that are less fractionated (Fig. 2c and Supplementary Fig. 5). In fact, the degree of fractionation bias is significantly correlated with the degree of diploidization (Fig. 2d). The dominant subgenome carries 56% of the 1*x* equivalent syntelog set (average of 5,293 sliding 100-gene windows), and every chromosome in the recessive subgenomes has at least 46 single-copy syntelogs (i.e., not detected in the other chromosomes) (Supplementary Fig. 6). Moreover, dominant chromosomes tend to be larger in assembled scaffold sizes (Fig. 2e). These results suggest that the establishment of dominant and recessive subgenomes played a crucial role in efficient gene fractionation and in the differentiation of chromosomal sizes.

In line with subgenome fractionation dominance, genes in the dominant subgenome tend to show higher expression levels compared with corresponding syntelogs in recessive subgenomes (Fig. 2f). Thus, the dominant/recessive distinction is clear in both gene retention and gene expression. It is noteworthy that, despite the clear chromosome-level expression dominance, 47% (972/2,055) of syntelog pairs showed higher expression levels in copies on the recessive subgenomes (Fig. 2g), showing a significant contribution to gene activity by these subgenomes as well.

The odd-numbered subgenomes of *Nepenthes* with clear subgenome dominance strongly suggest an allopolyploid history underlying this dominance pattern (Alger and Edger, 2020). Our attempts to use subgenome-specific *k*-mers identified by SubPhaser, which has successfully phased the subgenomes of many well-known allopolyploids (Jia et al., 2022), did not result in complete subgenome phasing in *Nepenthes* (Supplementary Fig. 7). Disentangling exact polyploidization history is sometimes difficult, especially in high-ploidy lineages (e.g., decaploid cotton (Wang et al., 2016)). Chromosome-scale syntenic comparisons between *Nepenthes* and related lineages, once they are available, will be necessary to discern among alternative scenarios.

### Sex chromosome evolution

Individual *Nepenthes* plants exclusively produce male or female flowers (dioecy), in contrast to the hermaphroditic flowers of their closest relatives and most other angiosperms (Fig. 1b). An XY-type sex determination system has independently evolved in this genus (Scharmann et al., 2019), but the sex chromosome has so far remained unidentified. To locate the Y-linked chromosomal region (YLR) in our genome assembly of a male *N. gracilis*, we mapped sequencing reads of double digest restriction-site associated DNA (ddRAD-seq) from wild, sex-identified *N. gracilis* individuals (Scharmann et al., 2019). The result showed a clear, sharply delimited signal of male-specific sequences in a 1 Mb region on chromosome 20 (Fig. 3a; Supplementary Fig. 8). Chromosome 20 is a member of one recessive subgenome (Fig. 2a), suggesting that the sex chromosome evolved from an ancestral chromosome that experienced relaxed purifying selection. Interestingly, the YLR has no homologous region on the corresponding X-chromosome in the female genome assembly (Supplementary Fig. 9), implying that it is fully hemizygous, similar to the independently evolved YLR of *Asparagus* (Harkess et al., 2020).

**Figure 3.**
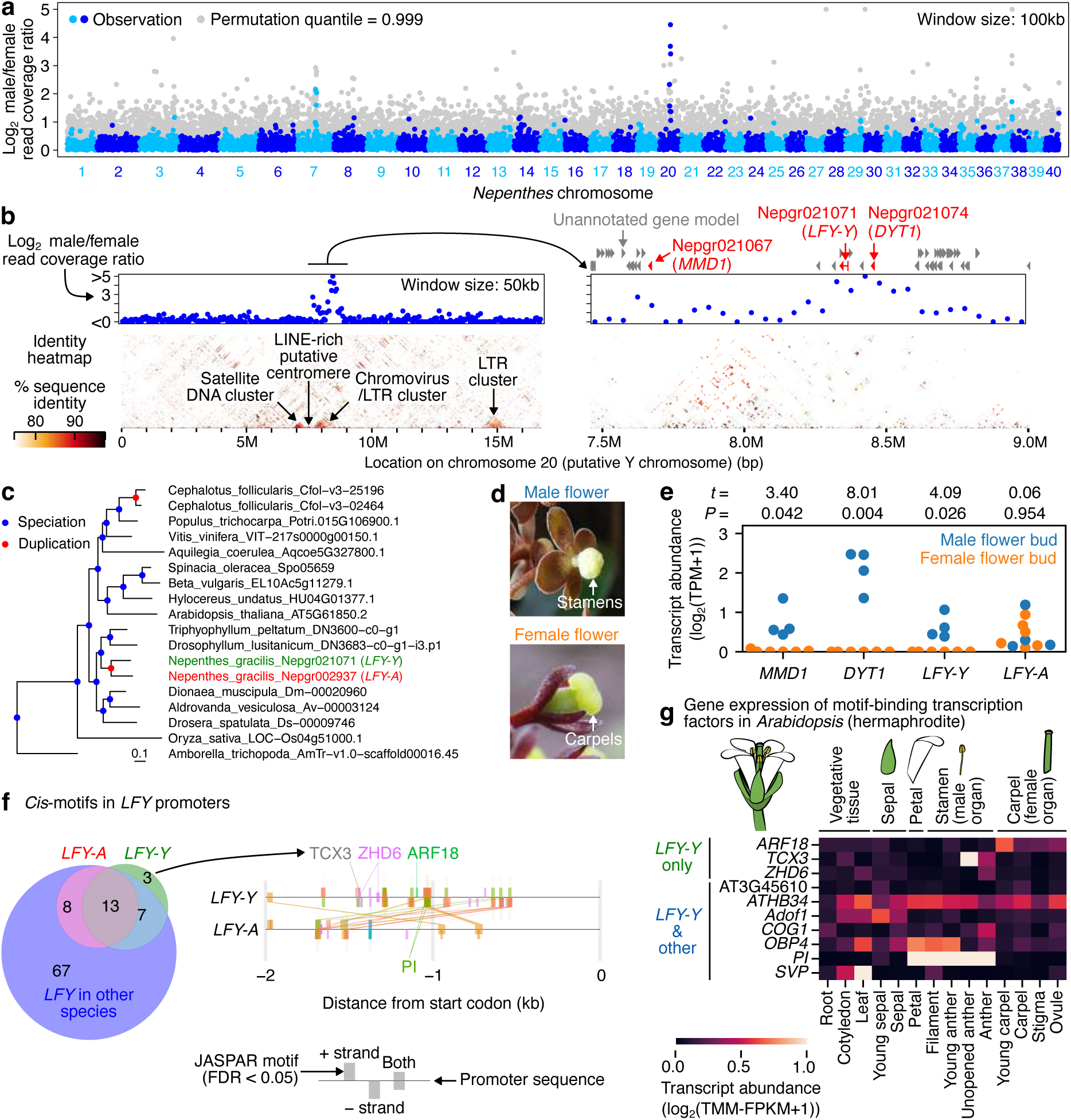
The male-specific chromosomal region harbors transcriptional regulators of flower development. (**a–b**) Analysis of genomic regions specific to male individuals in *N. gracilis*. ddRAD-seq reads from 11 males and ten females were mapped to the reference genome, and the male/female ratio is shown. In the bottom panels of **b**, tandem repeats are visualized with identity heatmaps using StainedGlass (Vollger et al., 2022). The range of the right panel corresponds to the range of the black line on the left panel. Positions of Trinotate-annotated and unannotated gene models are indicated in red and gray, respectively. The position of the putative centromere, which is typically LINE-rich in *N. gracilis*, is indicated in **b** (for details, see Supplementary Fig. 10). LINE, long interspersed nuclear elements; LTR, long terminal repeat. (**c**) Phylogenetic relationships of *LFY* genes in angiosperms. Duplicates in *Nepenthes* are indicated with colors. The bar indicates 0.1 substitutions per nucleotide site. Supplementary Fig. 13a provides a complete phylogeny. (**d**) Male and female flowers of *N. gracilis*. The male flower picture is licensed under CC BY-NC 4.0 (https://creativecommons.org/licenses/by-nc/4.0/) by HP Lim. (**e**) Expression of genes in the male-specific region. RNA-seq experiments were performed in cultivated individuals of *Nepenthes* spp. (Supplementary Table 4) and reads were mapped to the *N. gracilis* genome. *P* values and *t* statistics of Welch’s *t*-tests are provided above the plot. TPM: transcripts per million. (**f**) Promoter differentiation of duplicated *LFY* in *N. gracilis*. The transcription factor-binding motifs in JASPAR (FDR < 0.05) were detected in the 2-kb promoter sequences of all genes in **c**, and their overlap is shown in the Venn diagram. A comparison of the duplicated *LFY* promoters in *N. gracilis* is shown to the right. Commonly found motifs are connected with lines in order of proximity to the transcription start site. Three motifs specifically found in the *LFY-Y* promoter are indicated. (**g**) Tissue-specific expression patterns of transcription factor genes in *A. thaliana* flowers (Klepikova et al., 2016). The gene expression heatmap shows the genes encoding transcription factors binding to the motifs that are detected in *LFY-Y*. The flower illustration is licensed under CC BY 4.0 (https://creativecommons.org/licenses/by/4.0/) by Frédéric Bouché.

A major unresolved problem in sex chromosome research is the evolution of suppressed recombination. The lack of recombination throughout the *Nepenthes* YLR is implicit in the population-resequencing approach we used for the delimitation of the YLR. Although hemizygosity would in itself imply a lack of recombination, an alternative explanation is that the *Nepenthes* YLR is recombination-depauperate due to adjacency to the satellite-rich putative centromere (Fig. 3b and Supplementary Fig. 10). Pericentromeric regions, due to their heterochromatin and low recombination rates, might have favored the formation of Y-linked regions and other supergenes, as suggested in other plant and animal systems (Schwander et al., 2014; Li et al., 2015; Horiuchi et al., 2022; Potente et al., 2022; Akagi et al., 2023).

### Male-specific genes in the Y-linked chromosomal region

Notably, the male-specific region, which spans approximately 1 Mb in the 16.7-Mb chromosome, contains *DYSFUNCTIONAL TAPETUM 1* (*DYT1*), which is the only fully male-linked gene known to date in *Nepenthes* (Scharmann et al., 2019) (Fig. 3b). *DYT1* encodes a bHLH transcription factor that, in *Arabidopsis*, functions in the cell maturation of tapetum, the nutritive cell layer that aids microsporogenesis in the developing anther (Zhang et al., 2006). An ortholog of *MALE MEIOCYTE DEATH 1* (*MMD1*), which encodes a PHD-finger transcription factor whose loss causes male meiotic defects (Yang et al., 2003), is another transcription factor gene located in the male-specific region. While the characterized functions of *DYT1* and *MMD1* suggest their involvement in microspore development, which begins later in anther development, female flowers in *Nepenthes* not only lack microspores but also, almost entirely, staminal structures (Subramanyam and Narayana, 1971). We, therefore, hypothesized that another gene upstream of *DYT1* and *MMD1* locates to the male-specific region and determines floral sex. Consistent with this idea, we found a male-specific copy (*LFY-Y*) of the *LEAFY* (*LFY*) gene, which in hermaphroditic angiosperms encodes a plant-specific transcription factor that assigns the floral fate of meristems (Moyroud et al., 2010). *LFY* is one of the earliest expressed genes in flower development where it acts as a master regulator. Loss of *LFY* gene function converts lateral floral organs into leaf-like identity (Moyroud et al., 2010).

In *Nepenthes*, the three male-specific transcription factor genes (*DYT1*, *MMD1*, and *LFY-Y*) are embedded in a tandem cluster of long interspersed nuclear elements (LINEs), a group of retrotransposons (Fig. 3b and Supplementary Fig. 8). Transposable elements have not only accumulated in these intergenic regions but also in the introns of *LFY-Y* itself (Supplementary Fig. 11). It seems likely that this transposable element accumulation is associated with the suppressed recombination discussed above.

*LFY* is well known to be a highly conserved gene in angiosperms, where it is maintained as a single copy in most species (Moyroud et al., 2009). The *Nepenthes* genome harbors what is likely to be the principal *LFY* copy on an autosome (*LFY-A* on chromosome 3), which is not homeologous to the putative sex chromosome. Gene phylogeny (Fig. 3c), chromosomal syntenic groups (Fig. 2a), and the presence of introns (Supplementary Fig. 11) suggest that the two *LFY* copies emerged by a lineage-specific SSD rather than via WGD in *Nepenthes* after the split of the other carnivorous lineages in Caryophyllales, consistent with the emergence of dioecy in Nepenthaceae alone. This situation contrasts with *DYT1* and *MMD1*, both of which are maintained as single-copy genes in the decaploid genome of *Nepenthes*. As expected, *DYT1*, *MMD1*, and *LFY-Y* were expressed in the developing buds of male flowers but not in female buds (Fig. 3d,e). Furthermore, the male-specific expression of *LFY-Y* was maintained until flower maturation (Supplementary Fig. 12a). By contrast, we found no significant difference in *LFY-A* expression between male and female buds. Although this expression analysis has been performed by heterologous mapping to the *N. gracilis* genome using RNA-seq reads from related species (see Methods), the read mapping rates were comfortably high (75.9–83.9% in 31 samples), and we further confirmed by transcriptome assembly that the transcripts of *DYT1*, *MMD1*, and *LFY-Y* were detected only in male samples (Supplementary Fig. 12b–d). In the YLR, only these three genes could be functionally annotated with sequence similarity against the UniProt database (Supplementary Table 5), and they showed the highest male/female transcript abundance ratios in developing buds among all physically linked gene models in the region (Supplementary Fig. 12e). In summary, the three transcription factor genes in YLR could have been involved in establishing dioecy in *Nepenthes*.

### Neofunctionalization of the male-specific *LFY-Y* gene copy

To examine how two *LFY* copies acquired distinct expression patterns, we characterized *cis*-regulatory motifs in the promoters by mapping their motif sequences using the JASPAR database (Castro-Mondragon et al., 2022). With the false-discovery rate cutoff of 0.05, we detected 23 types of putative transcription factor-binding motifs in the *LFY-Y* promoters (Supplementary Table 7), of which three motifs were specifically found in *LFY-Y* among promoter sequences from other species: i.e., motifs bound by ARF18, TCX3, or ZHD6 (Fig. 3f). In *Arabidopsis*, *ARF18* is highly expressed in developing carpels (Fig. 3g) and encodes a transcriptional repressor (Liu et al., 2015), whereas TCX3 and ZHD6 are expressed mainly in anthers (Fig. 3g), suggesting roles for these regulators in modulating *LFY-Y* expression in both male and female organs.

These *LFY-Y*-specific motifs (ARF18, TCX3, and ZHD6) were detected 1.0-1.5 kb upstream from the start codon. This region may have been newly acquired by *LFY-Y* during sex chromosome evolution. Within this promoter region, we also detected paired PISTILLATA-binding (PI-binding) motifs only in *Nepenthes LFY-Y*, and interestingly, in *LFY* promoters of the comparator species spinach (*Spinacia oleracea*). LFY in fact also upregulates *PI* expression in *Arabidopsis* (Honma and Goto, 2000). PI is a B-class MADS-box protein that specifies petal and stamen identity in *Arabidopsis* and other plants (Theißen et al., 2016). Spinach evolved dioecy independently from *Nepenthes* (Fig. 1b), and the suppression of spinach *PI* converts male flowers into phenotypically normal female flowers (Sather et al., 2010), suggesting a potentially convergent mechanism underlying the evolution of dioecy among different Caryophyllales lineages through regulatory interactions between LFY, PI and other transcription factors. Taken together, these results suggest that *LFY-Y* was neofunctionalized in terms of expression pattern, likely as a consequence of the change in *cis*-regulatory elements. We detected five amino acid substitutions specifically found in LFY-Y in its C-terminus DNA-binding domain (Supplementary Fig. 13), suggesting potential changes in protein properties in addition to the expression patterns.

### Pitcher-tissue-specific prey capture responses

Besides dioecy, the trap leaf organization also serves as an evolutionary innovation in *Nepenthes*. A single *Nepenthes* leaf consists of segmented parts, including a carnivorous pitcher trap, which is further elaborated by tissue differentiations along the proximodistal axis, representing one of the most complex leaf shapes known among angiosperms (Tsukaya, 2014). In Caryophyllales, plant carnivory evolved before the origin of Nepenthaceae (Fig. 1b), presumably with the flypaper-type trapping mechanism and in a conventional leaf organization (Albert et al., 1992; Heubl et al., 2006; Renner and Specht, 2011), with the pitcher organization representing an evolutionary novelty that emerged in *Nepenthes*.

To examine whether gene expression patterns parallel unique pitcher tissue differentiation, we performed RNA-seq experiments in six dissected leaf parts: from proximal to distal, flat part, tendril, digestive zone, waxy zone, peristome, and lid (Fig. 4a–b). For a cross-species comparison, we also generated the pitcher tissue-specific transcriptomes of *Cephalotus*, which evolved pitcher leaves independently from *Nepenthes* in an altogether different angiosperm order (Fig. 1b), thus representing convergent evolution (Albert et al., 1992; Freund et al., 2022). We further integrated additional RNA-seq data from *Dionaea* (Bemm et al., 2016; Iosip et al., 2020; Procko et al., 2021) and *Arabidopsis* (Klepikova et al., 2016) to form a 4-species expression level dataset normalized with trimmed mean of M-values (TMM) (Robinson and Oshlack, 2010) for a total of 3,572 single-copy genes. Dimensionality reduction analysis with different methods consistently showed that, in gene expression profiles, the pitcher tissues of *Nepenthes* were distinct from conventional photosynthetic organs (Supplementary Fig. 14a). This contrasts with the pitcher leaves of *Cephalotus*, whose tissues showed expression profiles largely overlapping with the photosynthetic organs, suggesting that pitcher tissues in *Nepenthes* traps are much more strongly differentiated from ancestral photosynthetic leaf structures. This view is consistent with the contrasting patterns of the tissue-specific photosynthetic activity in the two pitcher plant lineages, with *Nepenthes* pitchers showing almost no photosynthetic assimilation (Pavlovič et al., 2007; Pavlovič, 2011).

**Figure 4.**
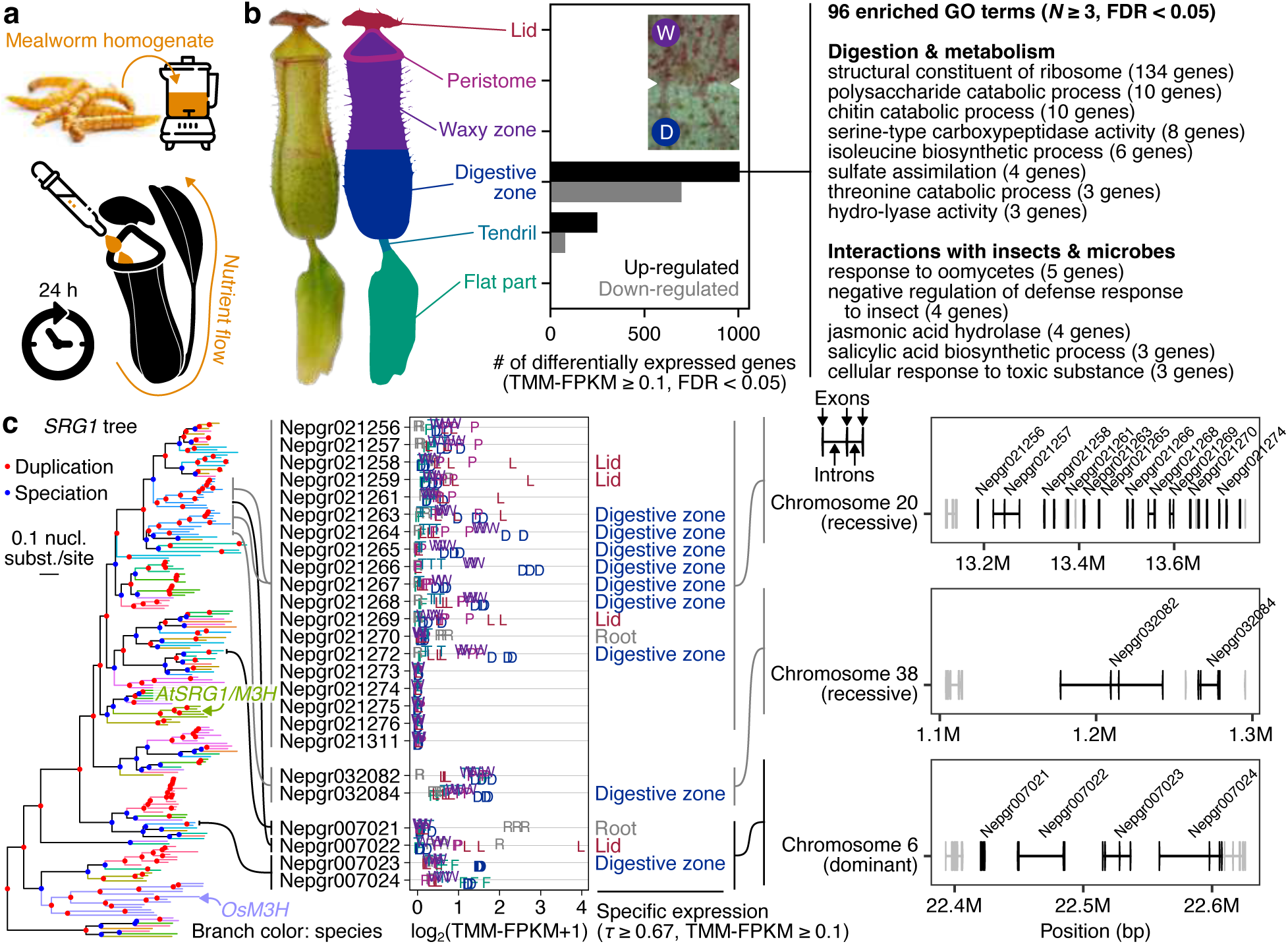
Pitcher tissue-specific gene expression. (**a**) The feeding experiment. (**b**) Prey-responsive genes in pitcher tissues. The boundary between the waxy zone and the digestive zone is shown as an inset. To the right, Gene Ontology terms enriched in the up-regulated genes in the digestive zone are shown. (**c**) The massive amplification of *SRG1* genes in *Nepenthes*. The maximum-likelihood phylogeny (left), expression level and tissue specificity (middle), and chromosomal location (right) are shown. The positions of *A. thaliana SRG1*/*M3H* and *Oryza sativa M3H* are indicated in the tree. The chromosomal locations of non-*SRG1* gene models are indicated in gray. Image credits: the mealworm photo and icons from freepik.com.

Next, we characterized tissue-specific prey capture responses in *Nepenthes*. Pitchers were treated with an insect homogenate to mimic prey capture, and tissues were harvested after 24 h with the expected onset of jasmonic acid-mediated (JA-mediated) prey responses (Yilamujiang et al., 2016). JA is a known regulator of gene expression in plants and can mediate plant-herbivore and pathogen interactions (Glazebrook, 2005; Erb et al., 2012), as well as prey-capture responses in the Caryophyllales lineage of carnivorous plants including *Nepenthes* (Pavlovič and Mithöfer, 2019). As anticipated, the number of significantly differentially expressed genes (DEGs, FDR < 0.05) was highest in the digestive zone, where digestive/absorptive glands (Freund et al., 2022) are directly exposed to the insect homogenate (Fig. 4b). A clear response of the digestive zone was further confirmed by dimensionality reduction analysis (Supplementary Fig. 14b). Significantly enriched Gene Ontology (GO) terms (Supplementary Table 8; Supplementary Table 9; Supplementary Table 10; Supplementary Table 11) among the up-regulated genes clearly indicated the activation of translation machinery upon prey capture (Supplementary Table 8), which would be anticipated for rapid response by the palette of necessary digestive enzymes. Other enriched terms include those presumably associated with prey digestion, nutrient metabolism, and biological interactions (Fig. 4b). The second-highest number of DEGs was found in the tendril, which connects the pitcher to the rest of the plant body. Interestingly, with the above threshold, we did not detect DEGs in other trapping leaf tissues, including the flat part, where absorbed nutrients flow through. These results suggest that prey response is well compartmentalized within a *Nepenthes* trapping leaf.

### Tandem gene clusters with pitcher-tissue-specific expression

The leaf complexity of *Nepenthes* may be reflected by considerable sub– or neofunctionalized duplicate gene expression in novel tissues, and amplification of gene copies is often mediated by tandem gene duplications (Durand and Hoberman, 2006). We therefore searched for tandem gene clusters with high tissue specificity in gene expression, measured by a metric called *τ* (Yanai et al., 2005), which outperformed alternative measures in a benchmark study (Kryuchkova-Mostacci and Robinson-Rechavi, 2016). A conspicuous example we detected was gene clusters of *SENESCENCE-RELATED GENE 1* (*SRG1*) orthologs (Callard et al., 1996). This senescence marker gene encodes cytoplasm-localized melatonin 3-hydroxylase (M3H) and produces cyclic 3-hydroxymelatonin, whose antioxidant activity is 15-fold higher than that of its precursor melatonin, and promotes growth in *Arabidopsis* and rice (Lee et al., 2016; Choi and Back, 2019; Lee and Back, 2022). This family of genes forms tandem duplicates on separate *Nepenthes* chromosomes within the same syntenic group (Fig. 4c), with the largest cluster (19 gene models with 15 cases with complete protein domain structure (Supplementary Fig. 15)) on chromosome 20 (the YLR-containing chromosome), which belongs to a recessive subgenome. Phylogenetic analyses suggested that the first copy of this cluster originated as a WGD duplicate whose counterpart in the dominant subgenome (chromosome 6) remains a single-copy gene (Nepgr007022). The tandem genes in the dominant subgenome showed well-conserved microsynteny among eudicots (Supplementary Fig. 15), suggesting ancient origins (>131 MYA, Fig. 1b). Interestingly, the tandem array on chromosome 20 contains many genes with expression specific to the digestive zone in the pitcher, which forms the interface to the digestive fluid, where peroxidases likely produce ROS to facilitate proteolysis for prey degradation (Chia et al., 2004; Hatano and Hamada, 2012; Fukushima et al., 2017; Wal et al., 2022). Given this characteristic expression, the SRG1 proteins may participate in scavenging cytotoxic reactive oxygen species (ROS) produced during prey digestion and nutrient absorption.

To further examine whether other carnivory-related genes similarly formed tandem clusters, we analyzed genes encoding digestive enzymes. Using the list of experimentally confirmed digestive fluid proteins, many of which are digestive enzymes (Fukushima et al., 2017), we characterized their phylogeny, synteny, and expression (Supplementary Fig. 16). Among the 11 families of genes we analyzed, we detected eight tandem gene clusters (six on recessive subgenomes) that were formed in *Nepenthes* after its split from close relatives in Caryophyllales. Tandem clusters encoding typical digestive enzymes (aspartic protease, class III peroxidase, glycoside hydrolase family 19 chitinase, purple acid phosphatase, RNase T2, and β-1,3-glucanase) included genes whose transcript abundance was highest in the digestive zone, with increased expression following the feeding treatment. These results suggest that specific tandem cluster evolution is not restricted to *SRG1* and may be found in many other genes involved in *Nepenthes* carnivory.

### Unequal subgenomic contributions for *Nepenthes*-specific genes

Since the male-specific region harboring *LFY-Y* and many tandem gene clusters expressed in the digestive zone (*SRG1* and digestive enzyme genes) were found in recessive subgenomes rather than the dominant subgenome, subgenomes may differentially serve as hosts of novel duplicated genes. However, these observations may also be explained by chance, since there is a 70% probability that a novel gene will occur in the recessive subgenome if the occurrence is proportional to the chromosome assembly size (528/753 Mb), and a 67.7% probability if it occurs proportionally to the number of genes (23,039/34,010 genes). Therefore, statistical analysis was necessary. To this end, we analyzed lineage-specific genes in the *Nepenthes* genome. First, DIAMOND BLASTP searches (Buchfink et al., 2021) were conducted against 20 other plant genomes to identify *Nepenthes* genes that are most similar to another *Nepenthes* gene rather than to genes from other species and thus likely to have emerged by gene duplication after the split from those other lineages. Identified lineage-specific genes were significantly enriched with those having specific expression in all analyzed tissues (Fig. 5b). Next, the types of duplication (i.e., WGD or SSD) and involved subgenomes (dominant or recessive) were estimated by whether the best-hit gene was on the same chromosome, on homeologous chromosomes, or on other chromosomes (Fig. 5a). Finally, the frequency of each category in tissue-specific genes was compared with the overall average obtained from all expressed genes. Interestingly, the recessive subgenomes tended to host more tissue-specific genes than the dominant subgenome. Such distribution shifts from the null expectation with all expressed genes were statistically significant in genes specifically expressed in six out of the seven analyzed tissues (Fig. 5b), including pitcher tissues, whose differentiation evolved in *Nepenthes* to form pitfall traps. The difference is likely to have emerged during functional divergence after gene duplications because the frequency of tandem duplications themselves was comparable between dominant and recessive subgenomes (Supplementary Fig. 17).

**Figure 5.**
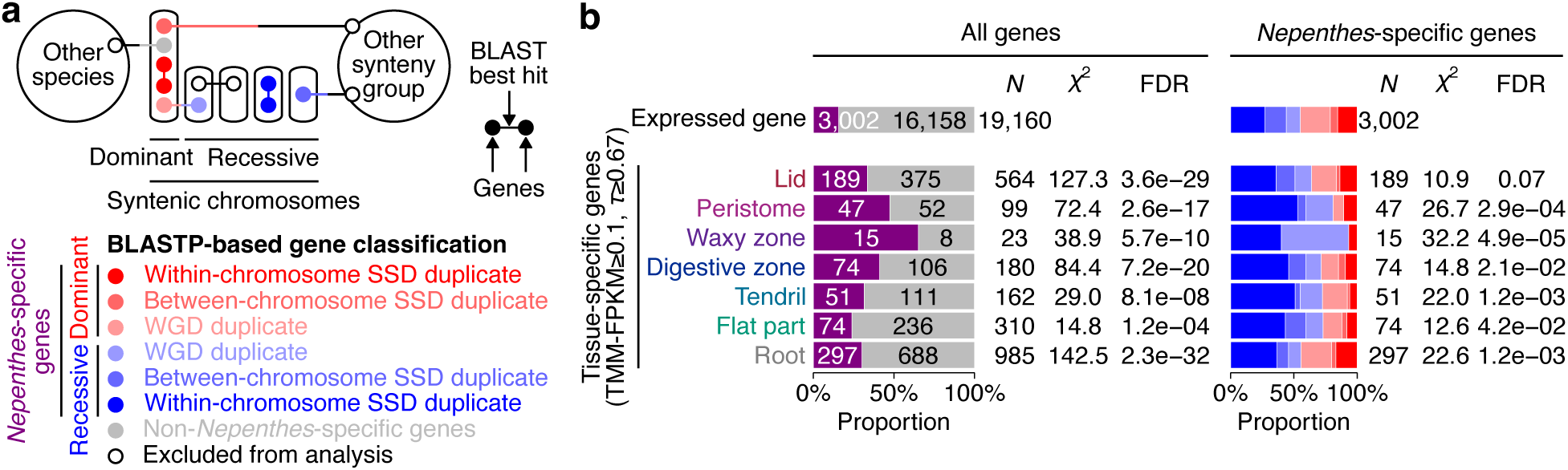
Differential subgenomic contributions to novel gene evolution. (**a**) Inference of duplication events in *Nepenthes*-specific genes. Duplication origins between small-scale duplications (SSDs) and whole-genome duplications (WGDs) are distinguished by the best hit of a DIAMOND BLASTP search (Buchfink et al., 2021). (**b**) Differential contributions of duplication categories in tissues-specific *Nepenthes* genes. Colors match those in **a**. The distributions of duplication categories in tissue-specific genes were compared with that of expressed genes using *χ*^2^ tests controlled for false discovery rates (FDR).

The increased contribution to the tissue-specific genes by recessive subgenomes was most pronounced among within-chromosome SSD duplicates, which are enriched in tandemly duplicated genes. The rate of protein evolution (*dN/dS*) was higher in SSD duplicates than in WGD duplicates, with the within-chromosome SSD duplicates showing the highest rates (Supplementary Fig. 18). Because there was no overall difference in *dN/dS* between dominant and recessive subgenomes, it is possible that subgenome dominance may have only transiently relaxed purifying selection in the recessive subgenomes, and that its signature is difficult to capture in the extant gene sequences, except for the substantial gene losses via fractionation discussed above (Fig. 2).

Different modes of gene duplication have been shown to exert different long-term impacts on the complements of genes retained in plant genomes. Generally, as similarly noted for many other plant systems (Freeling, 2009), WGD duplicates tend to be enriched for regulatory functions, whereas small-scale (e.g., tandem) duplicates are more likely to be enriched in functions related to plant defense (and possibly by extension, to plant carnivory). To examine how functional gene categories are differentially enriched in the dominant and recessive subgenomes of *Nepenthes*, we analyzed overrepresented GO categories (Supplementary Fig. 19, Supplementary Table 12, Supplementary Table 13, Supplementary Table 14, and Supplementary Table 15). Among other functions, the syntelogs on the dominant subgenome were significantly enriched for genes annotated with ‘flower development’, and ‘ethylene-activated signaling pathway’ GOs (Bonferroni-corrected *P* < 0.05). The syntelogs on the recessive subgenomes had potentially related GO enrichments such as ‘abaxial cell fate specification’, ‘floral meristem determinacy’, ‘leaf morphogenesis’, and various plant hormone GOs, including ‘response to jasmonic acid’. As such, the evolution of dominant and recessive subgenomes following polyploidization may have included both specialization and partitioning of ancestral regulatory networks, in a manner analogous to neofunctionalization and subfunctionalization, respectively, at the individual gene level (Conant and Wolfe, 2008). Regarding small-scale, local duplication events, dominant tandems were enriched in genes annotated with ‘cell surface receptor signaling pathway’ and ‘response to oomycetes’, whereas recessive tandems were enriched, for example, with GOs such as ‘methyl jasmonate methylesterase activity’ and ‘salicylic acid glucosyltransferase (glucoside-forming) activity’, which could conceivably be associated with prey recognition pathways (Hedrich and Fukushima, 2021).

Taken together, these results suggest that recessive subgenomes play an important role in hosting novel tissue-specific genes that evolved through SSDs, possibly in an environment of relaxed purifying selection compared to that in the dominant subgenome, and thereby to the unique biology of *Nepenthes*, including dioecy (Fig. 3) and carnivory (Fig. 4).

## Discussion

In this study, we elucidated the decaploid structure of the *Nepenthes* genome and identified a clear signature of 1:4 subgenome dominance (Fig. 2). We also highlighted how the four recessive subgenomes have contributed to evolutionary novelties in *Nepenthes* (Fig. 1), especially in relation to dioecy and carnivorous trapping leaves.

Our analysis of the organization of the male-specific chromosomal region (Fig. 3) suggested that a series of mutational events may have led to the evolution of dioecy in *Nepenthes* (Supplementary Fig. 20a). A likely scenario is an evolutionary path from hermaphroditism via gynodioecy towards full dioecy (Pannell and Jordan, 2022). First, an ancestral population of hermaphrodites could have given rise to a gynodioecious population through the loss-of-function of *MMD1* or *DYT1* in the ancestral X chromosome, which belongs to a recessive subgenome. This event would have resulted in the emergence of homozygous, recessive female-only plants without functional pollen. The linkage of *MMD1* and *DYT1* and the double loss-of-function on the X chromosome might have enhanced the female-only trait. Frequency-dependent selection may then have favored male function in hermaphroditic plants, until the gain of the masculinization gene on the ancestral Y chromosome. Such a masculinization gene should dominantly suppress the production of, or at least the function of, the carpels. From our association and expression analysis, *LFY-Y* was a plausible candidate for the masculinization gene, which, in the above scenario, would have resulted in the evolution of a completely dioecious proto-*Nepenthes* population. As such, our results prefer gynodioecy, rather than androdioecy, as a plausible intermediate step to full dioecy, because it is difficult to explain the linkage of *MMD1* and *DYT1* in the male-specific region if the masculinization gene evolved ahead of time to form androdioecious intermediates. Furthermore, *LFY-Y* expression in male flowers and *LFY-A* expression in both flower sexes are consistent with neofunctionalization in *LFY-Y* rather than *LFY* duplicate subfunctionalization (Conant and Wolfe, 2008) as a mode of functional change after gene duplication (Supplementary Fig. 20b). Indeed, the functional partitioning we describe here resembles the opposite of the sex-aggregating partial *LFY* redundancy and deletion scenario (Albert et al., 2002) to explain the evolution of the bisexual angiosperm flower from separate male and female reproductive axes controlled by two distinct *LFY* copies in gymnosperms.

In addition, our analysis of tissue-specific genes in *Nepenthes* pitcher leaves (Fig. 4) showed how molecular evolution likely paralleled the increased complexity of tissue organization in *Nepenthes*. As one of the most prominent examples, *SRG1* genes, which may participate in scavenging ROS during prey digestion and nutrient absorption, formed a massive tandem cluster in a recessive subgenome, and many of its members acquired tissue-specific expression in *Nepenthes*-specific tissues, such as the digestive zone. In contrast to the *LFY* duplication, the mode of functional divergence (i.e., neo– or subfunctionalization) in the *SRG1* cluster is not clear. However, it seems clear from gene expression and phylogenetic relationship data (Fig. 4c) that successive functional divergence has taken place during cluster formation. These instances of novel genes (i.e., *LFY-Y* and *SRG1*) occurred in recessive subgenomes, as do many other *Nepenthes*-specific genes that have acquired tissue-specific expression (Fig. 5), including digestive enzyme genes (Supplementary Fig. 16). This genome-wide trend (Fig. 5) suggests that WGDs influenced the fates of subsequently produced SSD duplicates through recessivity in a system with strongly divergent subgenome dominance patterns.

Our findings of novel gene duplicates suggest that the higher occurrence of tissue-specific genes after SSDs in recessive subgenomes may have contributed to adaptive potential in *Nepenthes*, and thus, to the maintenance of recessive subgenomes. Thus, recessive subgenomes have not degenerated completely, but have instead persisted for long periods of time (58.3–93.8 MY in *Nepenthes* (Scharmann et al., 2021)), contributing to the emergence of evolutionary novelties. Although the myriad of polyploids that have arisen in plant evolution may frequently represent evolutionary dead-ends (Van de Peer et al., 2017), but, as we have shown in *Nepenthes*, recessive subgenomes may have served as a source of adaptive potential in some highly radiated lineages. Our findings will therefore help revise models of karyotype stability and gene divergence during polyploid evolution.

## Methods

### Plant materials

For genome sequencing, male and female *Nepenthes gracilis* individuals were collected from the field. The collection date was 26 March 2019, and the collection locality was Tampines Avenue 10, N1°21’27.1” E103°55’49.9” (1.35753° N, 103.93053° E), Tampines, Singapore. The habitat was an open secondary forest on seasonally waterlogged flat ground with *Dillenia suffruticosa*, *Acrostichum aureum*, *Dicranopteris linearis*, sedges, and grasses. For transcriptome analysis of vegetative tissues, we purchased *N. gracilis* from CZ Plants (Ostrava-Radvanice a Bartovice, Czech Republic). The plants were grown on a mixture of sphagnum moss and perlite in the greenhouse. Since it was difficult to obtain fresh flower samples from *N. gracilis*, we obtained developing inflorescences from cultivars and other species (Supplementary Table 4). The inflorescence was dissected into early flower buds, late flower buds, young flowers, and mature flowers (Supplementary Fig. 21). Plants of *Ancistrocladus abbreviatus* (Ancistrocladaceae) (Bringmann et al., 1999) and *Drosophyllum lusitanicum* (Drosophyllaceae) were grown on soil in a greenhouse. Axenically grown strains of *Cephalotus follicularis* (Cephalotaceae) (Fukushima et al., 2017; Fukushima et al., 2021) and *Triphyophyllum peltatum* (Dioncophyllaceae) (Bringmann et al., 1990; Bringmann and Rischer, 2001) were maintained in the half-strength Murashige and Skoog solid medium (Murashige and Skoog, 1962) supplemented with 3% sucrose, 1× Gamborg’s vitamins, 0.1% 2-(N–morpholino)ethanesulfonic acid, 0.05% Plant Preservative Mixture (Plant Cell Technology) and 0.3% Phytagel, at 25°C (*C. follicularis*) or 20°C (*T. peltatum*) in continuous light.

### High-molecular-weight genomic DNA isolation

Young leaf tissues of male and female *Nepenthes gracilis* individuals from the wild were gathered, cleaned, and flash-frozen in liquid nitrogen, and then stored at –80°C prior to extraction. About 10 g of flash-frozen tissue was used for high-molecular-weight (HMW) genomic DNA isolation. The first step followed the BioNano NIBuffer nuclei isolation protocol, in which frozen leaf tissue was homogenized in liquid nitrogen, with a subsequent nuclei lysis step using IBTB buffer with spermine and spermidine added and filtered just before use. IBTB buffer consists of Isolation Buffer (IB; 15 mM Tris, 10 mM EDTA, 130 mM KCI, 20 mM NaCl, 8% (w/v) PVP-10, pH9.4) with 0.1% Triton X-100, and 7.5% (V/V) β-mercaptoethanol mixed in and chilled on ice. The mixture of homogenized leaf tissue and IBTB buffer was strained to remove undissolved plant tissue. 1% Triton X-100 was added to lyse the nuclei before centrifugation at 2000 ×g for 10 min to pellet the nuclei. Once the nuclei pellet was obtained, we proceeded with CTAB (cetyltrimethylammonium bromide) DNA extraction with modifications for the Oxford Nanopore sequencer as described previously (Michael et al., 2018). The quality and concentration of HMW genomic DNA was checked using Thermo Scientific™ NanoDrop™ Spectrophotometer, as well as on agarose gel electrophoresis following standard protocols. The genomic DNA was further purified with a Qiagen® Genomic-tip 500/G according to the protocol provided by the developer.

### Genome sequencing

HMW DNA for a male and a female plant was used to generate a sequencing library for use with Oxford Nanopore Technology (ONT) (SQK-LSK109; Oxford, England). The resulting libraries were run on a PromethION sequencer running for 48 h. All bases were called on the PromethION using Guppy v2.0, and the resulting fastq files were used for genome assembly.

### Genome assembly and polishing

The fastq sequencing data was filtered for >10-kb reads using seqtk v1.2 (https://github.com/lh3/seqtk). The filtered reads were assembled using WTDBG2 v2.2 (wtdbg2 – t64 –p19 –AS2 –e2 –L10000) (https://github.com/ruanjue/wtdbg2). A consensus genome assembly was generated by mapping reads >10 kb to the assembly with minimap2 v0.2 (https://github.com/lh3/minimap2), and then running Racon v1.3.1 (https://github.com/isovic/racon); the consensus process was repeated three times. The contig assembly was further polished using paired– end 2×150 Illumina sequence by first aligning the reads to the consensus assembly using minimap2 followed by running the assembly tool Pilon v1.18 (https://github.com/broadinstitute/pilon) three times using the paired-end 2×150 Illumina sequence data. Purge Haplotigs v1.0.0 (https://bitbucket.org/mroachawri/purge_haplotigs/src/master/) was applied to both the female and male Nanopore assemblies separately. The raw Illumina reads were aligned to the genome assemblies using bwa mem v0.7.17 (https://github.com/lh3/bwa), and input files were prepared using SAMtools v1.3 (https://github.com/samtools/samtools). Purge Haplotigs was then run using the prepared bam files and genome assembly.

### Hi-C scaffolding and syntenic path assembly

While Illumina-polished ONT-based wtdbg2 assemblies were generated independently for male and female specimens, we further improved the contiguity of the male assembly using HiRise scaffolding of Chicago and Dovetail Hi-C libraries by Dovetail Genomics (CA, USA) (Putnam et al., 2016) (Supplementary Fig. 22; Supplementary Table 1). Heterologous Hi-C scaffolding of the female genome assembly was performed using the female wtdbg2 assembly and the male Hi-C sequencing data. A list of Hi-C contact positions for the female was generated with Juicer v1.6 (https://github.com/aidenlab/juicer). This file was then used as input for 3D-DNA v170123 (https://github.com/aidenlab/3d-dna) to order and orient fragments of the genome assembly. Because of the lack of detectable synteny to any regions in the male assembly, an unnaturally large scaffold of the female Hi-C assembly was collapsed to its original contigs. We also employed another approach to scaffolding the female ONT assembly. With the male Hi-C assembly as a reference, we applied the syntenic path assembly to the female genome using the SynMap function of CoGe (Haug-Baltzell et al., 2017). Assembly statistics are available in Supplementary Table 2. The scaffold numbering in the final assembly corresponds to the chromosome numbers discussed in this paper.

### Feeding experiment

Dried mealworms (Batch No. L400518, MultiFit Tiernahrungs GmbH) were ground into a fine powder with a mortar and a pestle to prepare a 100 mg/ml homogenate in water. The homogenate was subsequently centrifuged at 2,000 ×g for 30 sec, and the supernatant (mealworm extract) was obtained. Upon the feeding experiment, we measured the total volume of digestive fluid and added the mealworm extract adjusted to 10% of the total volume to ensure a comparable concentration among pitchers with different sizes. The pitchers were then sealed using parafilm and left in the greenhouse for 24 h. The same procedure was applied to control plants fed with water alone.

### RNA extraction and transcriptome sequencing

The leaves of *N. gracilis* were dissected into the lid, peristome, waxy zone, digestive zone, tendril, and flat part (Supplementary Fig. 21). Digestive fluid was discarded. After washing with water and drying with a paper towel, the tissues were immediately frozen in liquid nitrogen. Tissues from five leaves (from multiple individuals) were pooled for one replicate, and a total of three biological replicates were prepared. Root samples were collected from untreated plants, and each replicate was derived from a single individual. Frozen samples were homogenized in liquid nitrogen using mortar and pestle. RNA extraction was performed with PureLink Plant RNA Reagent solution (Thermo Fisher) according to the manufacturer’s instructions. The RNA pellet was resuspended in RNase-free water at 4°C. After centrifuging at 14,000 rpm for 10 min at 4°C, the solution was transferred to a new tube and purified using an RNeasy Mini kit (Qiagen). While the above extraction method was used for *Nepenthes*, *Ancistrocladus*, *Triphyophyllum*, and *Cephalotus* (Supplementary Fig. 23) as well as leaves and roots of *Drosophyllum*, a modified CTAB protocol was employed for *Drosophyllum* flowers. The frozen *Drosophyllum* flower samples were homogenized with mortar and pestle. The ground sample was transferred to a pre-cooled 2-ml tube, and 0.75 ml of preheated 2×CTAB buffer (65°C) was added. The tube was shaken vigorously by using a vortex mixer and incubated in a thermoshaker at 1,400 rpm for 20 min at 60°C. An aliquot of 0.75 ml chloroform:isoamyl alcohol (25:1) was added and mixed vigorously. The homogenate was centrifuged at 15,000 rpm for 15 min at room temperature. The supernatant was then transferred to a new 2-ml tube. For RNA precipitation, 1.5 ml of 100% ethanol was mixed with 60 µl of 3 M aqueous sodium acetate and added to the supernatant, which was then shaken at 500 rpm for 60 min at room temperature. The homogenate was centrifuged at 15,000 rpm for 20 min at room temperature. The supernatant was discarded carefully so as not to disturb the pellet. The pellet was then washed with 1 ml of 75% ethanol. The supernatant was discarded, and the pellet was air-dried for 15 min in a thermoshaker at 37°C. The pellet was resuspended in 100 µl of RNase-free water at 4°C by shaking at 500 rpm for 15 min. The samples were again centrifuged at 15,000 rpm for 5 min at 4°C. The solution was transferred to a new tube, and the RNA was purified using the RNeasy Mini kit (Qiagen). The quality of RNA was examined using Nanodrop (Thermo Fisher) and the Qubit IQ assay kit (Invitrogen). Total RNA was sent to Novogene UK, where paired-end mRNA sequencing libraries were prepared using the Novogene NGS RNA Library Prep Set (PT042). Briefly, after the poly-A enrichment and fragmentation, the first strand cDNA was synthesized using random hexamer primers followed by the second strand cDNA synthesis, end repair, A-tailing, adapter ligation, size selection for 250–300-bp insert size, amplification, and purification. Libraries were paired-end sequenced for 150 bps with an Illumina Novaseq 6000 platform.

### Transcriptome assembly

Transcriptome assemblies for *A. abbreviatus*, *D. lusitanicum,* and *T. peltatum* were generated with the RNA-seq reads (Supplementary Table 4). The reads were preprocessed with fastp v0.20.0 (https://github.com/OpenGene/fastp) and assembled using Trinity (see Supplementary Table 6 for version) (https://github.com/trinityrnaseq/trinityrnaseq). Open reading frames (ORFs) were obtained with TransDecoder v5.5.0 with a minimum length of 50 bp (-m 50) (https://github.com/TransDecoder/TransDecoder). The longest ORFs among isoforms were extracted with the ‘aggregate’ function of CDSKIT v0.9.1 (https://github.com/kfuku52/cdskit).

### Repeat identification and annotation

For subsequent gene model prediction, repetitive elements on the reference genome were masked using RepeatMasker v4.0.9 (https://github.com/rmhubley/RepeatMasker) with a species-specific repetitive sequence library generated by RepeatModeler v2.0 (https://github.com/Dfam-consortium/RepeatModeler) (Supplementary Table 3). For further identification and annotation of repetitive elements, we used EDTA v2.1.0 (https://github.com/oushujun/EDTA). In order to identify the overall repetitiveness of genomes, we performed *de novo* repeat discovery with RepeatExplorer2 (Novák et al., 2020). We used a repeat library obtained from the RepeatExplorer2 analysis of Illumina paired-end reads. All clusters representing at least 0.005% of the genomes were manually checked, and the automated annotation was corrected if needed. Contigs from the annotated clusters were used to build a repeat library. Transposable element protein domains (Neumann et al., 2019) found in the assembled genomes were annotated using the DANTE tool available from the RepeatExplorer2 Galaxy portal (https://galaxy-elixir.cerit-sc.cz/). Further repeat density distribution plots shown in Supplementary Fig. 10 were made using shinyCircos (https://venyao.xyz/shinycircos/) and pyGenomeTracks v3.6 (Lopez-Delisle et al., 2021).

### Analysis of tandem repeat clusters

We used StainedGlass v0.4 (https://github.com/mrvollger/StainedGlass) to identify repeat sequence clusters. The genomic distribution of repeat sequences was visualized using HiGlass v0.10.1 (https://github.com/higlass/higlass) and HiGlass Manage (https://github.com/higlass/higlass-manage) with the gene annotation track (https://github.com/higlass/gene_annotations).

### Transcriptome assembly for gene modelling

The transcriptome assembly was performed using a combination of genome-guided and *de novo* approaches. The genome-guided approach employed StringTie v2.1.4 (https://github.com/gpertea/stringtie) with aligned reads from HiSat2 v2.1.0 (https://github.com/DaehwanKimLab/hisat2). For the *de novo* approach, we first ran Trinity v2.8.5 with default parameters, followed by running TransAbyss v2.0.1 (https://github.com/bcgsc/transabyss) for multiple *k*-mers (51, 61, 71, 81, 91, and 101). The resulting files from both approaches were merged to generate a single high-confidence transcriptome assembly using EvidentialGene v2022.01.20 (https://sourceforge.net/projects/evidentialgene/). This approach was repeated for six RNA-seq libraries (DRR461683–DRR461688).

### Gene model prediction

Gene model prediction was performed using a combination of *ab initio* and homology-based approaches. First, six transcriptome assemblies were splice-aligned against the genome using PASA v2.3.3 (https://github.com/PASApipeline/PASApipeline). The longest ORFs from these PASA alignments were also extracted using TransDecoder v.5.5.0. Next, we employed the *ab initio* gene prediction tool GeneMark-ES v4.65 (Brůna et al., 2020) and Braker v2.1.2 (https://github.com/Gaius-Augustus/BRAKER) to produce two separate sets of candidate gene models on the reference genome soft-masked by RepeatMasker as described above. The initial RNA-seq alignments for Braker were produced using STAR aligner v2.7.2b (https://github.com/alexdobin/STAR). The final prediction step in Braker was carried out using Augustus v3.3.2 (https://github.com/Gaius-Augustus/Augustus). Braker was run for six RNA-seq libraries. Homology-based predictor GeMoMa v1.6.1 (Keilwagen et al., 2018) was used to produce two additional sets of candidate gene models using gene models from *Arabidopsis thaliana* (Athaliana_167_TAIR10 from Phytozome) and *Populus trichocarpa* (Ptrichocarpa_444_v3.1 from Phytozome). All candidate gene models were then combined to form a single high-quality set of 34,010 gene models using EVidenceModeler v1.1.1 (https://github.com/EVidenceModeler/EVidenceModeler). The completeness of gene models was evaluated with BUSCO v5.3.2 (https://gitlab.com/ezlab/busco). The male gene models were transferred to the female genome assembly using GeMoMa.

### Gene annotation

Functional annotation of predicted coding sequences was performed with Trinotate v3.2.1 (https://github.com/Trinotate/Trinotate) with the DIAMOND BLASTP search (v2.0.15, E-value cutoff = 0.01, https://github.com/bbuchfink/diamond) against the UniProt database downloaded on June 21, 2022, resulting in the Gene Ontology annotation in 53% (18,037/34,010) of gene models. SignalP v4.1 (Petersen et al., 2011) and TMHMM v2.0c (Krogh et al., 2001) were used to predict signal peptides and transmembrane domains, respectively. Coding sequences were used for RPS-BLAST v2.9.0 searches (Camacho et al., 2009) against Pfam-A families (El-Gebali et al., 2019) (released on November 25, 2020) with an E-value cutoff of 0.01 to obtain protein domain architectures.

### Analysis of synteny and whole-genome duplication

Syntelogs between homologous *Nepenthes* chromosomes were identified using JCVI v1.2.7 (https://github.com/tanghaibao/jcvi) with the MCscan pipeline. The identification of syntelogs between species was performed using SynMap2 (https://genomevolution.org/wiki/index.php/SynMap2), which internally uses LAST for sequence alignments (Kiełbasa et al., 2011), and then fractionation bias was analyzed with FractBias (Joyce et al., 2017). The reproducible links are as follows: *Vitis* versus *Nepenthes* (https://genomevolution.org/r/1myic) and *Vitis* versus *Coffea* (https://genomevolution.org/r/1myu9). Synonymous divergence of paralogous pairs was obtained using WGDdetector v1.1 (Yang et al., 2019).

### Analysis of the male-specific region

The male-specific region was delimited on the male genome assembly using re-sequencing data (ddRAD-seq, (Peterson et al., 2012)) of 11 male and ten female individuals sampled from wild populations, including data from previous work (Scharmann et al., 2019). Male versus female read coverage was evaluated by mapping the read data to the genome with bwa mem v0.7.17-r1188, and alignments were filtered with SAMtools v1.12 against non-primary and supplementary alignments, and in the case of paired-end reads, enforcing the “properly-paired” status. Read depth per sample was counted in genomic windows using bedtools v2.30.0 (https://github.com/arq5x/bedtools2), and compared between the sexes by taking the log_2_ of the ratio of male and female normalized depth sums given by

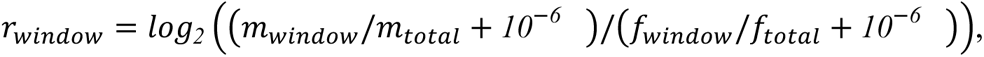

where *m* is the sum of male read counts and *f* is the sum of female read counts. Window-specific null distributions for this statistic were obtained by 1,000 permutations of the sex assignment of the 21 individuals. Male-specific sequences were further evaluated by counting *k*-mers (*k* = *16*) in the ddRAD-seq data of males and females using KMC v3.1.0 (https://github.com/refresh-bio/KMC), requiring *k*-mers to occur at least two times. Observed *k*-mers were classified as male-specific if they were present in at least nine out of 11 male samples and absent in all ten female samples, using kmc_tools. All possible alignments of such male-specific *k*-mers in the reference genome were found by bwa mem, allowing at most one mismatch. The abundance of such *k*-mer alignments was counted in genomic windows using bedtools. Window-specific null distributions for the abundance of male-specific *k*-mers were generated by 1,000 permutations of the sexes and repetition of the above kmc_tools and alignment procedure.

### Analysis of differential expression and GO enrichment

RNA-seq reads were preprocessed, pseudo-aligned to reference, and TMM-corrected using AMALGKIT v0.6.8.0 (https://github.com/kfuku52/amalgkit), which internally uses fastp, kallisto v0.48.0 (https://github.com/pachterlab/kallisto), and edgeR v3.36.0 (Robinson et al., 2010). OrthoFinder-based single-copy genes were used for the cross-species TMM correction of expression levels in *A. thaliana* (Brassicaceae), *C. follicularis*, *D. muscipula* (Droseraceae), and *N. gracilis*. In *N. gracilis*, the gene models of the male assembly were used for all samples, including flower samples from other species and cultivars. One flower sample (DRR461757) was removed from the analysis due to a low mapping rate (4.7%, Supplementary Table 4). Differentially expressed genes between fed and unfed samples were detected using edgeR.

### Analysis of *cis*-regulatory elements

The 2-kb sequences upstream of the start codons of *LFY* genes were obtained with SeqKit v2.3.1 (https://github.com/shenwei356/seqkit). The JASPAR CORE v2022 non-redundant set of experimentally defined transcription factor binding sites for plants (Castro-Mondragon et al., 2022) was used to search *cis*-regulatory elements using FIMO in the MEME Suite v5.4.1 (Bailey et al., 2015) with the false discovery rate (FDR) cutoff value of 0.05.

### Species tree inference

A total of 1,614 single-copy genes conserved in land plants were searched in the genomes and transcriptomes from the 20 species using BUSCO with the Embryophyta dataset in OrthoDB v10 (embryophyta_odb10) (Kriventseva et al., 2019). All genes marked as single-copy (S) or fragmented (F) were extracted, while those marked as duplicated (D) or missing (M) were treated as missing data (Supplementary Fig. 24). In-frame codon alignments were created by aligning translated protein sequences with MAFFT v7.475 (Katoh and Standley, 2013), trimming by ClipKIT v1.3.0 (https://github.com/JLSteenwyk/ClipKIT), and back-translation by CDSKIT v0.10.2 (https://github.com/kfuku52/cdskit). For each single-copy gene, nucleotide and protein maximum-likelihood (ML) trees were generated using IQ-TREE v2.2.0.3 (https://github.com/iqtree/iqtree2) with the GTR+R4 model and the LG+R4 model, respectively. The collection of 1,614 single-copy gene trees was subjected to the coalescence-based species tree inference with ASTRAL v.5.7.3 (https://github.com/smirarab/ASTRAL). Additionally, concatenated alignments were generated with catfasta2phyml v2018-09-28 (https://github.com/nylander/catfasta2phyml) and used as input of IQ-TREE for the nucleotide and protein ML tree inference with the above substitution models. *Amborella trichopoda* was used as the outgroup for rooting.

### Divergence time estimation

The divergence time estimation was performed with, as input, the ML species tree and the concatenated codon alignment of single-copy genes. Fossil constraints used in previous studies (Zhang et al., 2017; Suetsugu et al., 2023) were introduced with NWKIT v0.11.2 (https://github.com/kfuku52/nwkit). Species divergence was estimated with MCMCTREE in the PAML package v4.9 (https://github.com/abacus-gene/paml). Priors and parameters were chosen as described in the tutorial (http://abacus.gene.ucl.ac.uk/software/paml.html). Branch lengths and substitution model parameters were pre-estimated using BASEML with a global clock using the GTR+G model.

### Orthogroup classification

Gene sets from the 20 species (Supplementary Table 6) were grouped into 126,597 orthogroups by OrthoFinder v2.5.4 (https://github.com/davidemms/OrthoFinder) with the inferred species tree as a guide tree. For downstream analysis, we extracted 12,123 orthogroups with genes from at least 50% of species and no more than 1,000 genes (Supplementary Table 16).

### Gene tree inference

For each orthogroup, the nucleotide ML tree was generated with the GTR+G4 model as described above. The confidence of tree topology was evaluated with ultrafast bootstrapping (--ufboot 1000) with the optimization by hill-climbing nearest neighbor interchange (-bnni). The ML tree was used as the starting gene tree for GeneRax v2.0.4 (https://github.com/BenoitMorel/GeneRax) to generate a rooted, species-tree-aware gene tree. The divergence times of gene trees were inferred by the reconciliation-assisted divergence time estimation using RADTE v0.2.0 (https://github.com/kfuku52/RADTE), which uses a dated species tree as a reference to anchor the divergence time of gene tree nodes (Fukushima and Pollock, 2020).

### Analysis of gene trees

Branching events in gene trees were categorized into speciation or gene duplication by a species-overlap method (Huerta-Cepas et al., 2007). The *dN/dS* values were obtained for all branches in the 12,123 orthogroup trees using the *mapdNdS* approach with mapnh v2 (Guéguen and Duret, 2018) with parameter estimation using IQ-TREE according to a previous report (Fukushima and Pollock, 2020). Nonsynonymous codon substitutions in *LFY-Y* were estimated with IQ-TREE and were mapped to the protein structure with CSUBST v.1.1.0 (https://github.com/kfuku52/csubst) (Fukushima and Pollock, 2023).

### Data visualization

Phylogenetic trees were visualized using the R package ggtree v3.2.0 (https://github.com/YuLab-SMU/ggtree). General data visualization was performed with Python packages matplotlib v3.6.1 (https://github.com/matplotlib/matplotlib) and seaborn v0.12.0 (https://github.com/mwaskom/seaborn) as well as the R package ggplot2 v3.3.5 (https://github.com/tidyverse/ggplot2). The protein structures were visualized using Open-Source PyMOL v2.4.0 (https://github.com/schrodinger/pymol-open-source).

## Data Availability

Raw data and results are available at Dryad (https://doi.org/10.5061/dryad.xsj3tx9mj). The *N. gracilis* genome assembly and gene models are available from the DNA Data Bank of Japan (DDBJ) with the accession numbers BSYO01000001 to BSYO01000176 The *N. gracilis* genome assemblies are also available on CoGe (https://genomevolution.org/coge/) (genome ID: male assembly, 61566; female Hi-C assembly, 61892; female syntenic path assembly, 61931) and Dryad (https://doi.org/10.5061/dryad.xsj3tx9mj). DNA and mRNA sequencing reads were deposited to DDBJ (PRJDB15224, PRJDB15738, PRJDB15742, and PRJDB15737) and EBI (PRJEB20488), and the accession numbers are shown in Supplementary Table 1 and Supplementary Table 4.

## Code Availability

Scripts used in this study are available at Dryad (https://doi.org/10.5061/dryad.xsj3tx9mj).

## Acknowledgments

We acknowledge the following sources for funding: the Sofja Kovalevskaja programme of the Alexander von Humboldt Foundation (K.F.), a Human Frontier Science Program (HFSP) Young Investigators Grant RGY0082/2021 (K.F. and T.R.), Deutsche Forschungsgemeinschaft (DFG) Research Grants (454506241 to K.F., 415282803 to R.H., and 699/14-2 to G.B.), JSPS KAKENHI JP20H05909 (K.S.), United States National Science Foundation grants (1442190 to V.A.A. and 2030871 to T.R. and V.A.A.), United States Department of Agriculture-National Institutes of Food and Agriculture grant 2019-67012-37587 (K.J.G.), and research-leave funding from the School of Biological Sciences, Nanyang Technological University to V.A.A. and C.L. that supported *Nepenthes gracilis* collecting and sequencing. Matti Niissalo, Jia Hui Ang, Qiu Yan Tan, and Gillian Khew (Singapore Botanic Gardens, National Parks Board, Singapore) are thanked for their assistance with collecting and processing the *Nepenthes gracilis* material for long-read DNA sequencing. We thank Joachim Danz, Hirofumi Doi, and Leo Steffen for providing additional plant materials of *Nepenthes*, and Traud Winkelmann for propagating *in-vitro* cultures of *Triphyophyllum peltatum*. Computations were partially performed on the National Institute of Genetics (NIG) supercomputer, the Data Integration and Analysis Facility at the National Institute for Basic Biology, the Erlangen National High Performance Computing Center, the University at Buffalo Center for Computational Research, and High-Performance Computing Clusters at the University of Würzburg.

## Author Contributions

V.A.A. and K.F. conceptualized the project. F.S., G.V., G.B., Y.W.L., C.L., V.A.A., and K.F. acquired materials. Y.W.L., C.L., T.P.M., S.M., and K.S. conducted and/or oversaw DNA extraction and sequencing. F.S., E.C., T.P.M., V.A.A., and K.F. conducted genome assembly. F.S., L.C., M.F., T.W., and K.F. conducted RNA extraction. S.R. conducted gene model prediction. F.S., A.M., M.R., M.S., V.A.A., and K.F. analyzed the results. F.S., M.S., V.A.A., and K.F. wrote the manuscript with input from all authors. K.J.G. and T.R. contributed to improving the manuscript. D.B., R.H., V.A.A., and K.F. supervised the project and coordinated collaborations. All authors reviewed the manuscript.

## Supplementary Information for

Subgenome dominance shapes novel gene evolution in the decaploid pitcher plant *Nepenthes gracilis* Franziska Saul^#^, Mathias Scharmann^#^, Takanori Wakatake, Sitaram Rajaraman, André Marques, Matthias Freund, Gerhard Bringmann, Louisa Channon, Dirk Becker, Emily Carrol, Yee Wen Low, Charlotte Lindqvist, Kadeem J. Gilbert, Tanya Renner, Sachiko Masuda, Michaela Richter, Gerd Vogg, Ken Shirasu, Todd P. Michael, Rainer Hedrich, Victor A. Albert*, and Kenji Fukushima*

## List of Supplementary Materials

Supplementary Methods

Supplementary Text 1

Supplementary Tables 1–18 (separate file)

Supplementary Figs. 1–24

Supplementary References

Supplementary Dataset (separate file on Dryad: https://doi.org/10.5061/dryad.xsj3tx9mj)

## Supplementary Texts

### Supplementary Text 1. No strong gene-tree signal for inter-lineage allopolyploidization

Previous phylogenetic studies placed Nepenthaceae sister either to the Droseraceae (Brockington et al., 2009; Schäferhoff et al., 2010; Yang et al., 2018; Yao et al., 2019) or to the ADD clade (Meimberg et al., 2000; Cameron et al., 2002; Renner and Specht, 2011; Walker et al., 2017; Walker et al., 2018). This may be explained by the possibility that Nepenthaceae emerged by the allopolyploidization between two lineages: one close to Droseraceae and another close to the ADD clade. To test this possibility, we analyzed the 17 eudicot species (Supplementary Table 6) using Grampa v1.3.1 (https://github.com/gwct/grampa), which accounts for individual gene tree topology to distinguish auto– and allopolyploidy events (Thomas et al., 2017). To obtain a high-confidence set of gene trees, including lineage-specific gene duplication events that provide the signal of WGDs, we extracted BUSCO genes with the eudicot dataset of OrthoDB v10 (eudicots_odb10), including those labeled as duplicated. Using the translated sequences aligned with MAFFT and TrimAl, the ML trees were reconstructed using IQ-TREE with the LG+R4 substitution model and were rooted by NOTUNG (Chen et al., 2000). The analysis of the 2,326 gene trees using Grampa with *N. gracilis* assigned as H1 yielded the best multi-labeled tree which infers an allopolyploidization between the ancestral lineages of Droseraceae and the ADD clade, with 21 mapped duplication events that were judged as supporting the sister relationship between *N. gracilis* gene(s) and Droseraceae gene(s) (Supplementary Table 17). However, manual inspection of the allopolyploidization-supporting 24 gene trees that contain multiple *N. gracilis* genes revealed prevalent *N. gracilis*-specific duplications and paralogy or misplacement of *N. gracilis* genes. As a result, no analyzed gene trees actually showed the topology expected from the allopolyploidization scenario: i.e., ((*N. gracilis* gene 1, ADD genes), (*N. gracilis* gene 1, Droseraceae genes)).

Another aspect of the data that could potentially indicate a history of allopolyploidization is the topologies of phylogenies for genes that have returned to single-copy after the event. Here, we took advantage of the advanced diploidization which has occurred since polyploidization events took place in *Nepenthes*: the set of BUSCO genes (eudicots_odb10) is largely single-copy in *Nepenthes* (Supplementary Fig. 3d). If *Nepenthes* involved allopolyploid hybridization between the ADD and Droseraceae lineages followed by re-diploidization, we would expect that single-copy genes place *Nepenthes* either as sister to the Droseraceae or sister to the ADD families (and their relative proportions possibly manifesting as subgenome dominance, see main text). Strong gene tree incongruence of this kind is indeed observed in the data (quartet support only ∼0.45 in the ASTRAL species tree, Supplementary Fig. 3e). However, gene tree incongruence will not necessarily derive from hybridization but can also be caused by incomplete lineage sorting (ILS): i.e., rapid and nearly simultaneous speciation events among the three lineages (Nepenthaceae, Droseraceae, and ADD families). To distinguish between these alternatives, we applied Species Networks applying Quartets (SNaQ) (Solís-Lemus and Ané, 2016), an ILS-aware (coalescent) method to infer phylogenetic networks on protein sequences of single-copy genes in *Nepenthes*. Since the inference of phylogenetic networks is computationally intensive, the number of taxa on a gene tree must be kept relatively small. We, therefore, pruned the available gene trees to subsets of nine taxa, and in the case that taxa were represented by more than one gene copy, these taxa were discarded entirely (treated as missing data; consistent with the method used for species tree inference in the main text; Supplementary Fig. 24). We evaluated ten taxon subsets including the same seven ingroup taxa (*Nepenthes*, *Triphyophyllum*, *Drosophyllum*, *Ancistrocladus*, *Dionaea*, *Aldrovanda*, and *Drosera*), and two randomly chosen outgroup taxa. Starting from a bifurcating topology inferred by ASTRAL v5.7.8 (Zhang et al., 2018), phylogenetic networks with zero and then one hybrid edge were inferred and optimized using the SNaQ approach with PhyloNetworks v0.15.3 (https://github.com/crsl4/PhyloNetworks.jl) with 24 parallel attempts. While network optimization with one hybrid edge by SNaQ was successful in all ten taxon sets, *Nepenthes* did not descend from a hybridization event in any of them. Instead, it was usually the Droseraceae lineage which was constructed as hybridized with a ghost lineage directly descending from the ingroup stem (seven taxon sets), or the ADD clade as descending from such an event (two taxon sets), or the ADD lineage as hybridized with the *Nepenthes* lineage (one taxon set). Although these analyses do not support the hypothesis that *Nepenthes* is an allopolyploid hybrid involving ancient Droseraceae and/or the ancient ADD clade, it must be noted that there is as yet no satisfying explanation for the observed gene tree incongruence. In particular, neither the bifurcating species trees nor the optimized phylogenetic networks with one hybrid edge do fit the observed gene trees well (QuartetNetworkGoodnessFit v0.4.0 (Cai and Ané, 2021), all empirical *P* values < 10^-5^).

To be thorough, we analyzed gene location on subgenomes. If allopolyploidy occurred between the two distant lineages, genes on particular subgenomes should show the signal of particular phylogenetic affinity. A total of 33.4% (129/386) of the gene trees with the allopolyploidization signal (including those containing only one *N. gracilis* gene) are located on the dominant subgenome, which is comparable and not significantly different from the expectation with all genes (32.3%, 10,971/22,621, 418 gene models are on unanchored scaffolds and are thus excluded; *χ*^2^ test, *P* = 0.79, *χ*^2^ = 0.069). Therefore, our results do not support the possibility of the allopolyploidization events between the two distant lineages (i.e., ancestral Droseraceae and ancestral ADD clade). Still, there are phylogenetic signals that support the affinity of *N. gracilis* genes with Droseraceae, and this may be explained by ancient hybridization events before polyploidization. In this scenario, to explain the subgenomic frequencies of gene trees supporting alternative topologies, enough time would have been needed for allelic recombination to occur after hybridization and before polyploidization. Homeologous exchange (Bird et al., 2018), if it did occur, would also contribute to gene shuffling between subgenomes.

Supplementary Tables

Supplementary Table 1. DNA sequencing.

Supplementary Table 2. Genome assembly statistics.

Supplementary Table 3. Repetitive elements in the male assembly annotated by RepeatMasker.

Supplementary Table 4. RNA sequencing.

Supplementary Table 5. Annotation and expression of *Nepenthes* genes.

● gene_id: gene ID. One gene per line.
● orthogroup: Orthofinder-based orthogroup ID to which the gene is assigned.
● sprot_best: UniProt ID of DIAMOND BLASTP best hit.
● sprot_alias: Alias ID (if any) of sprot_best.
● sprot_coverage: DIAMOND BLASTP hit coverage. Q and H indicate the coordinates in queries and hits, respectively.
● sprot_identity: The percentage of sequence identity in sprot_coverage.
● sprot_evalue: DIAMOND BLASTP E-value of sprot_best.
● sprot_recnam: Gene name of sprot_best.
● signalp_start: Signal peptide start coordinate predicted by SignalP.
● signalp_start: Signal peptide end coordinate predicted by SignalP.
● signalp_score: SignalP score.
● tmhmm_expa: Expected number of amino acids in transmembrane helices predicted by TMHMM.
● tmhmm_predhel: Number of predicted transmembrane helices by N-best.
● tmhmm_topology: Topology predicted by N-best. i = inner, o = outer.
● kegg_gene: KEGG gene ID matched to sprot_best.
● kegg_orthology: KEGG orthology ID of kegg_gene.
● go_ids: GO IDs annotated to sprot_best.
● go_aspects: ordered GO aspects of go_ids.
● go_terms: ordered GO terms of go_ids.
● busco_id: BUSCO gene ID.
● busco_status: BUSCO status.
● busco_sequence: BUSCO hit coordinate of gene_id.
● busco_score: BUSCO score.
● busco_length: BUSCO gene length.
● busco_orthodb_url: OrthoDB URL of the BUSCO gene.
● busco_description: Gene description of the BUSCO gene.
● feature_size: Size of coding sequences (bp).
● num_intron: Number of introns.
● chromosome: Chromosome, scaffold, or contig on which the gene locates.
● start: Start coordinate of the gene.
● end: End coordinate of the gene.
● strand: Strand of the gene.
● TISSUE_REPLICATE: Expression level (Log_2_ TMM-FPKM).
● is_enough_tmmfpkm: Whether the maximum TMM-FPKM value is greater or equal to 0.1.
● max_tissue: Tissue with the maximum value of average TMM-FPKM.
● tau: Yanai’s tau for the tissue specificity of gene expression obtained from TMM-FPKM values.
● is_enough_tau: Whether the tau value is greater or equal to 0.67.
● fdr_TISSUE: False discovery rate of the TISSUE versus TISSUEFed comparison by edgeR.
● is_enough_tissue_tmmfpkm_TISSUE: Whether the TMM-FPKM value is greater or equal to 0.1 in the TISSUE or TISSUEFed samples.
● logFC_TISSUE: log_2_ fold change values (TISSUEFed/TISSUE) estimated by edgeR.
● logCPM_TISSUE: Log_2_ average of CPM values over compared samples (TISSUE and TISSUEFed).
● F_TISSUE: F statistics of the TISSUE versus TISSUEFed comparison by edgeR.
● PValue_TISSUE: Uncorrected *P* value of the TISSUE versus TISSUEFed comparison by edgeR.

Supplementary Table 6. Plant genomes for evolutionary analyses.

Supplementary Table 7. JASPAR motifs detected in LFY promoters.

Supplementary Table 8. GO enrichment analysis of the genes up-regulated in the digestive zone.

Supplementary Table 9. GO enrichment analysis of the genes down-regulated in the digestive zone.

Supplementary Table 10. GO enrichment analysis of the genes up-regulated in the tendril.

Supplementary Table 11. GO enrichment analysis of the genes down-regulated in the tendril.

Supplementary Table 12. GO enrichment analysis of the syntelogs on the dominant subgenome.

Supplementary Table 13. GO enrichment analysis of the syntelogs on the recessive subgenomes.

Supplementary Table 14. GO enrichment analysis of the tandem duplicates on the dominant subgenomes.

Supplementary Table 15. GO enrichment analysis of the tandem duplicates on the recessive subgenomes.

Supplementary Table 16. The number of genes and codon alignment statistics in individual orthogroups.

Supplementary Table 17. Detected gene duplication mapping to the best multi-labeled species tree.

Supplementary Table 18. Identification of BLAST-based orthologs encoding digestive fluid proteins.

## Supplementary Figures

**Supplementary Fig. 1.**
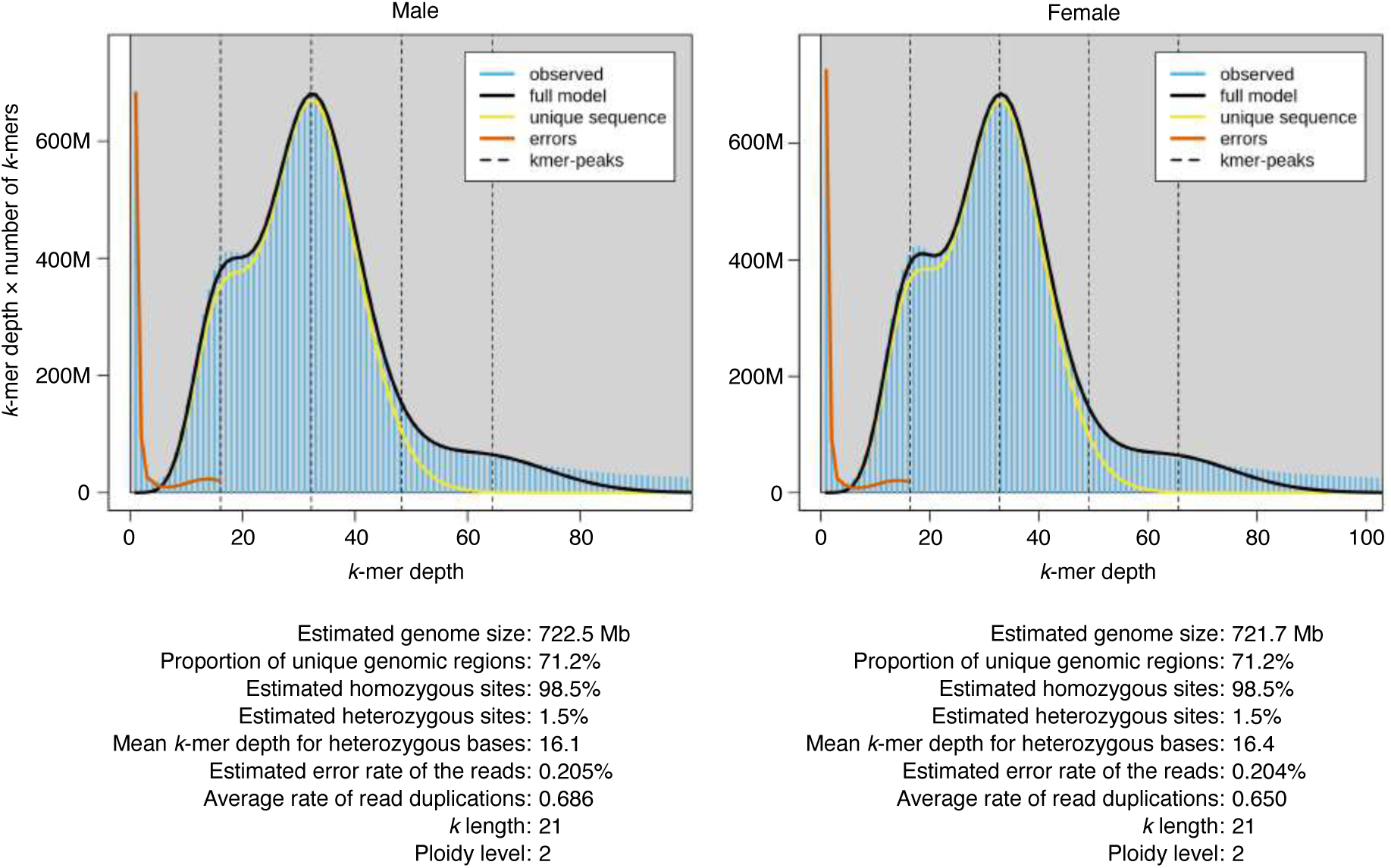
The *k*-mer frequency distribution of the *Nepenthes gracilis* male and female individuals. Illumina short reads (Supplementary Table 1) were analyzed using GenomeScope v2.0 (https://github.com/tbenavi1/genomescope2.0) (Ranallo-Benavidez et al., 2020).

**Supplementary Fig. 2.**
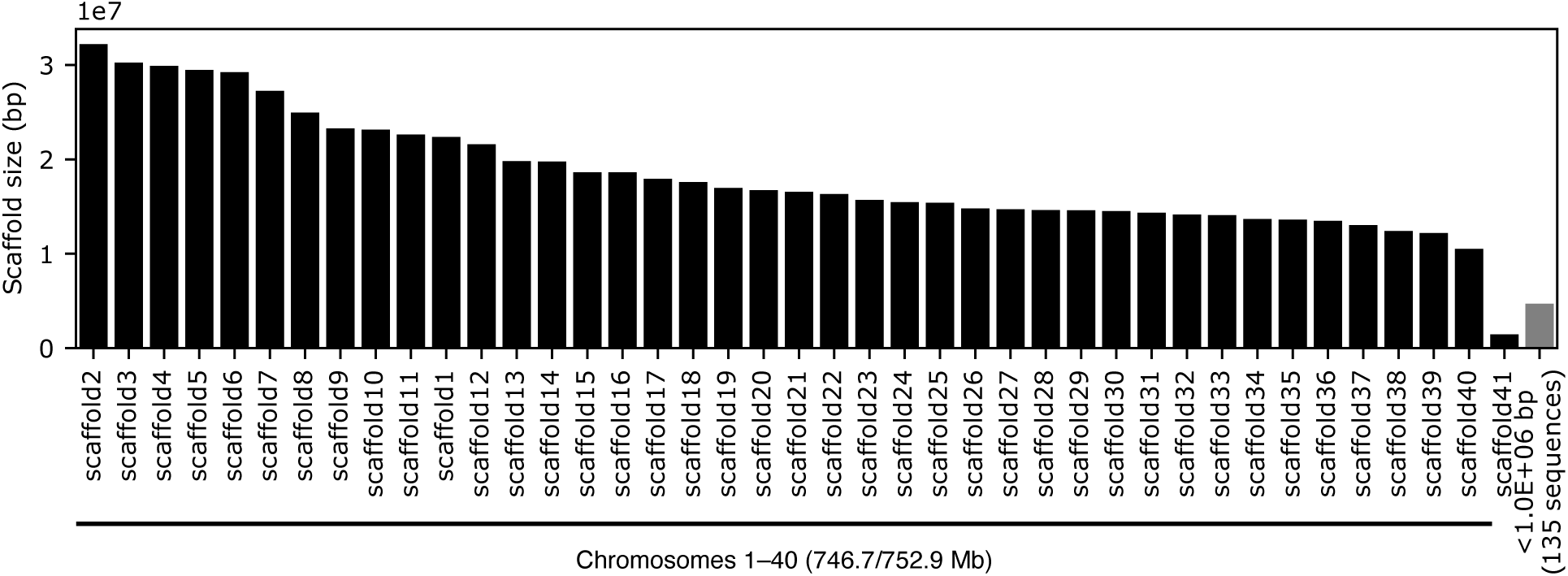
The scaffold size distribution of the *Nepenthes gracilis* male genome 1414 assembly. Scaffolds smaller than 1 Mb are merged to the right.

**Supplementary Fig. 3.**
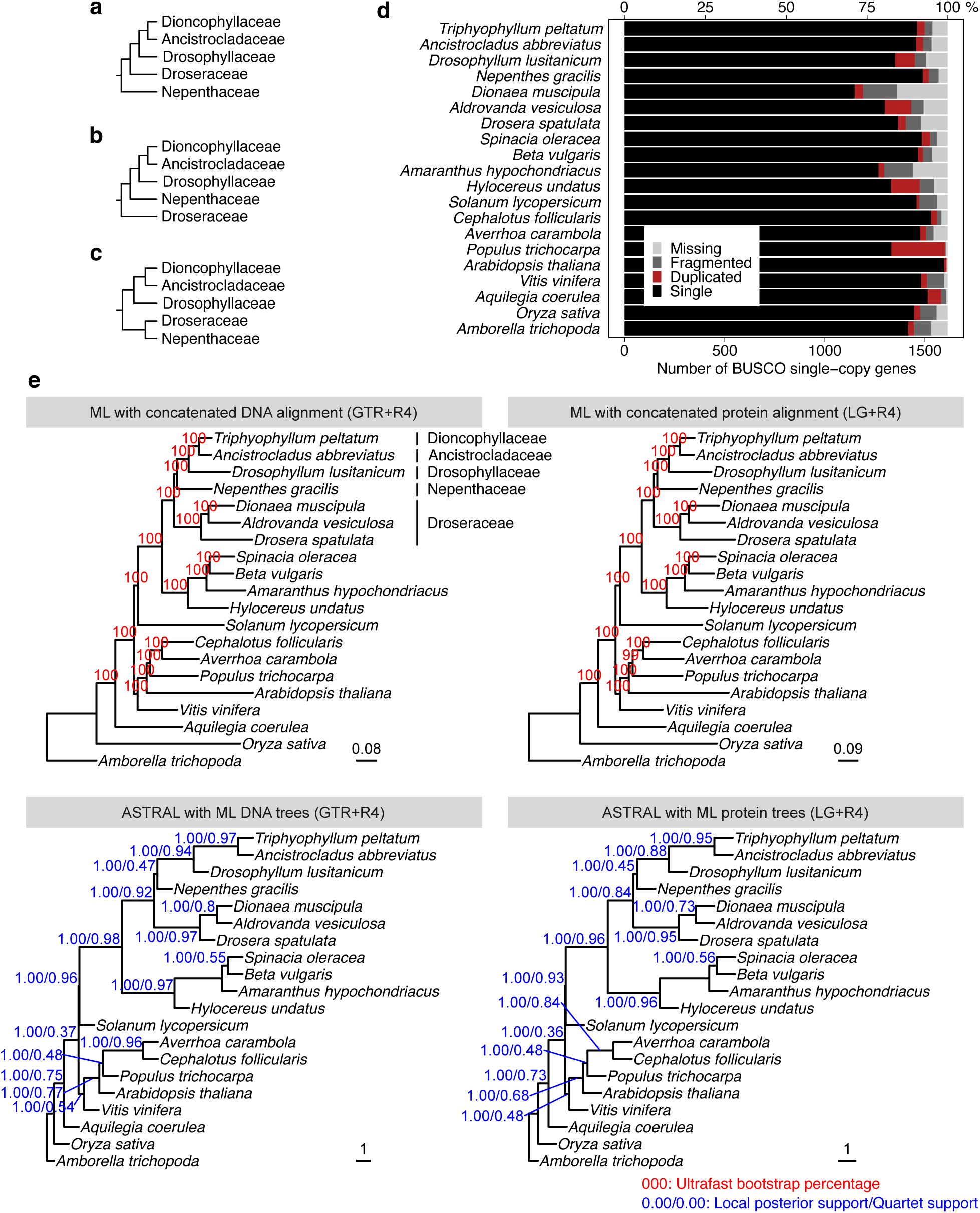
Phylogenetic hypotheses on the position of Nepenthaceae. (**a-c**) The positions in **a** (Lledo et al., 1998; Crawley and Hilu, 2012a; Crawley and Hilu, 2012b), **b** (Meimberg et al., 2000; Cameron et al., 2002; Renner and Specht, 2011; Walker et al., 2017; Walker et al., 2018), and **c** (Brockington et al., 2009; Schäferhoff et al., 2010; Yang et al., 2018; Yao et al., 2019) are supported by previous research. (**d**) Phylogenetic relationships inferred by the maximum-likelihood (ML) method and a coalescence-based method (ASTRAL) with DNA and protein alignments of 1,614 single-copy genes conserved in Embryophyta (defined by BUSCO embryophyta_odb10; 2,575,635 nucleotide sites). All analyses support the tree topology shown in **b**. Branch lengths correspond to nucleotide and amino acid substitutions per site, respectively, in the upper left and the upper right panel. In the lower panels, lengths of internal branches correspond to coalescent units, while those of terminal branches are not estimated (see https://github.com/smirarab/ASTRAL/blob/master/astral-tutorial.md) and are arbitrarily shown only for visualization purposes.

**Supplementary Fig. 4.**
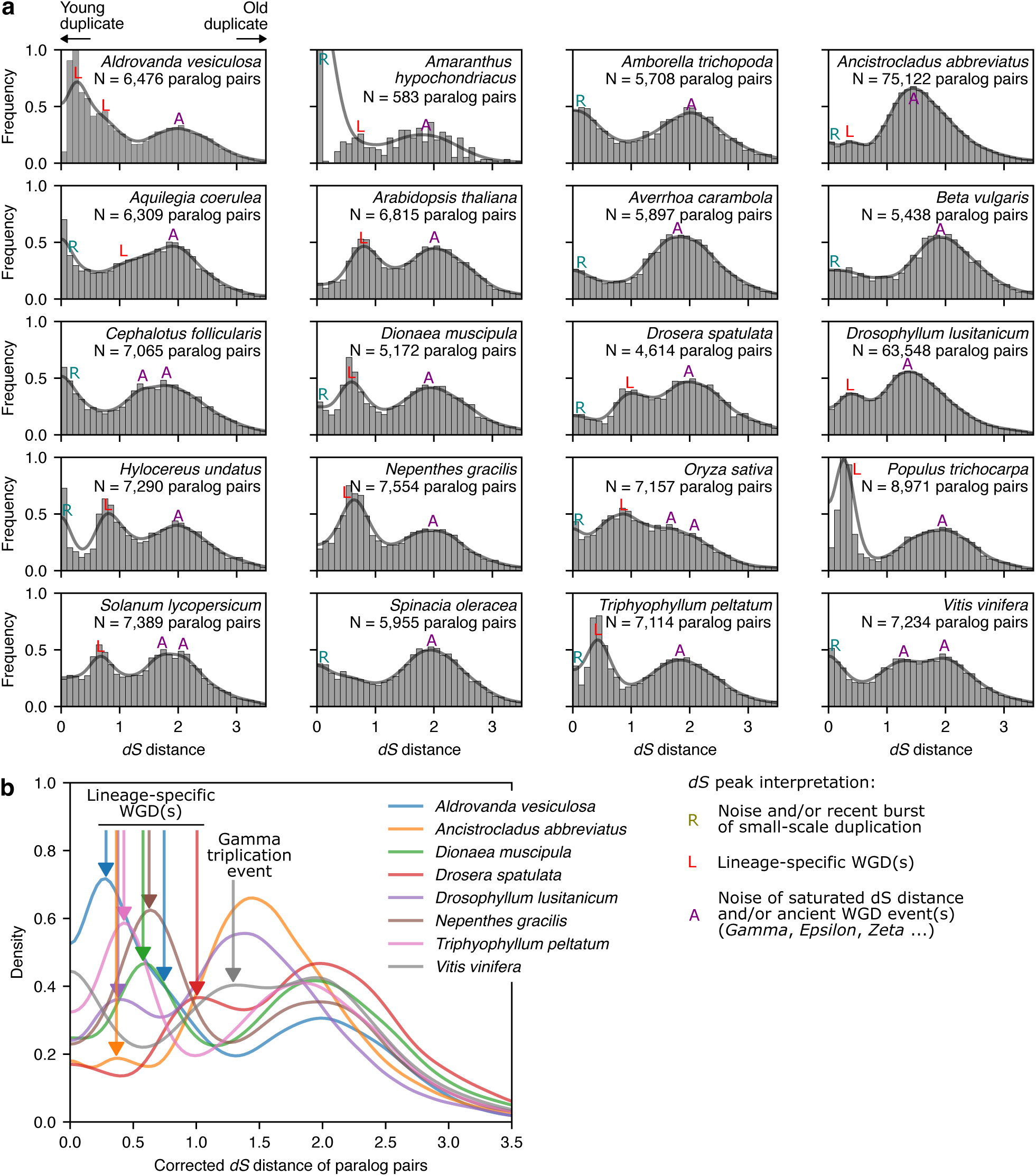
Distribution of synonymous divergence (*dS*) in paralogous gene pairs within the genome. (**a**) Analysis of the 20-genome dataset with WGDdetector (Yang et al., 2019). Histograms are overlaid with a kernel density plot with the reflective lower boundary. Our interpretation of detected *dS* peaks is labeled in the plot. Recent bursts of small-scale gene duplications (Zwaenepoel et al., 2019), as well as errors, such as incompletely purged haplotigs and splicing variants, can produce a peak near *dS* = 0 (R). Saturated *dS* distance and/or ancient WGD events, such as the *gamma* hexaploidization event shared by all extant eudicots, may produce a peak with large *dS* values (A). Lineage-specific WGDs (L) are expected to be located between the R and A peaks. (**b**) A comparison of lineage-specific WGDs in carnivorous Caryophyllales species. *V. vinifera* is included to anchor the location of the *gamma* hexaploidization event.

**Supplementary Fig. 5.**
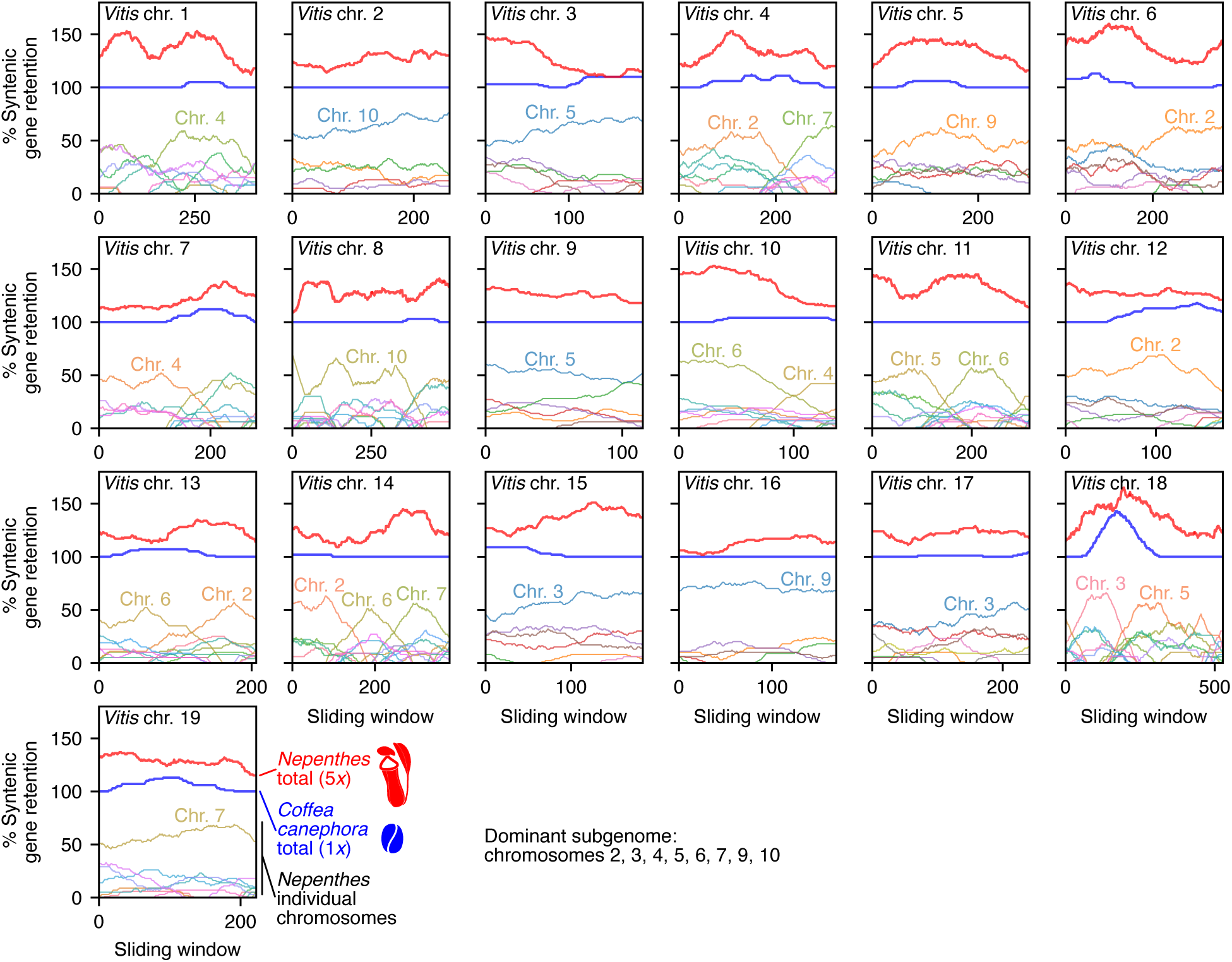
Analysis of fractionation bias in the *Nepenthes* genome. The rate of syntenic gene retention is evaluated by aligning the *Nepenthes* genome (5*x*) against the *V. vinifera* genome (1*x*), which is known not to have experienced a WGD after the *gamma* hexaploidization event (Jaillon et al., 2007). Analysis of the *C. canephora* genome (1*x*) with the same FractBias parameters (Joyce et al., 2017) is shown for comparison. Dominant chromosomes with clear signals are indicated in the plot. The results are reproducible on CoGe with the following link: *V. vinifera – N. gracilis* (https://genomevolution.org/r/1myic) and *V. vinifera – C. canephora* (https://genomevolution.org/r/1n0tj).

**Supplementary Fig. 6.**
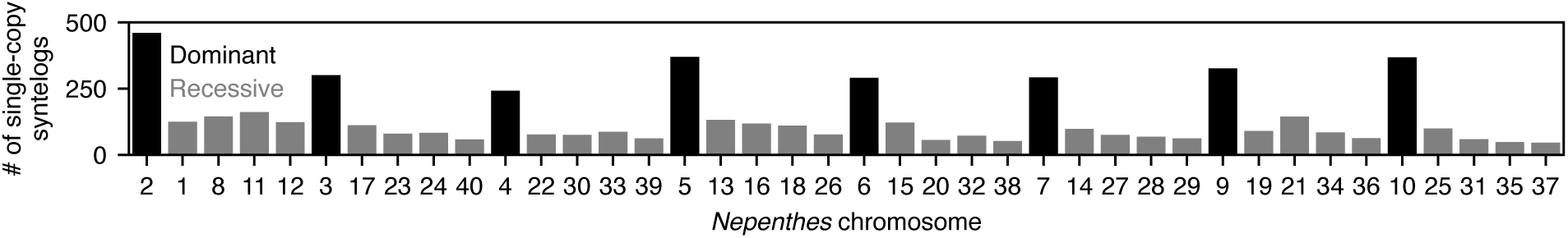
**Chromosome-wise number of single-copy syntelogs against the *Vitis* genome in the *Nepenthes gracilis* genome.**

**Supplementary Fig. 7.**
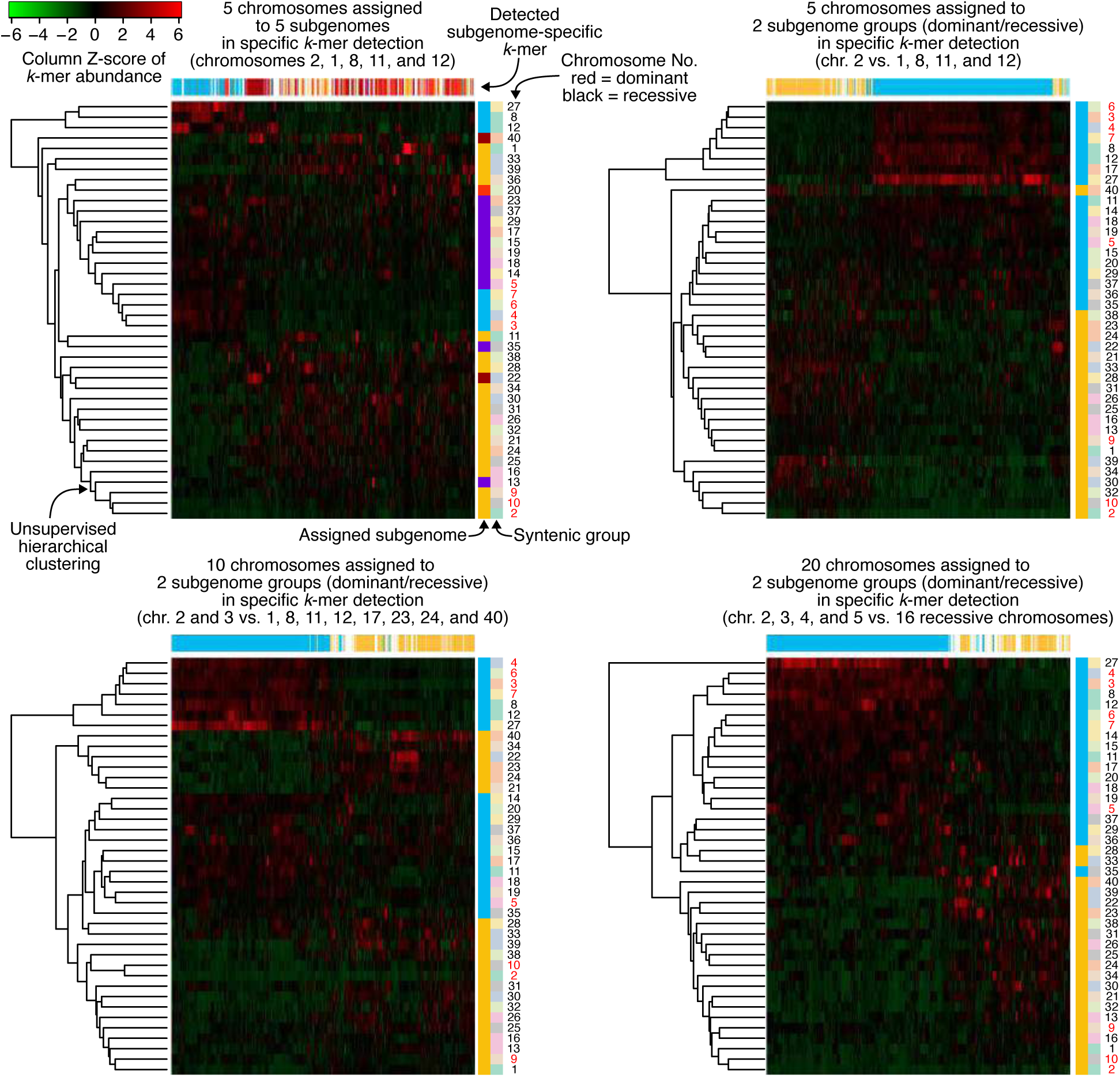
Subgenome phasing with specific *k*-mers. The *Nepenthes* male genome was analyzed using SubPhaser v1.2.5 (https://github.com/zhangrengang/SubPhaser) (Jia et al., 2022) with different sets of chromosomes in the subgenome specification for the subgenome-specific *k*-mer detection. Heatmaps show the relative abundance of differential *k*-mers. Syntenic groups of homologous chromosomes are shown according to Fig. 2a. The subgenome phasing worked partially, with the robust clustering of several chromosomes in the dominant subgenome (i.e., the clade of chromosomes 3, 4, 6, and 7 and the clade of chromosomes 2 and 10), but was not perfect even though different parameter settings were employed. No subgenome-specific *k*-mer was detected with all dominant and recessive chromosomes (8 and 32, respectively) specified in the subgenome-specific *k*-mer detection.

**Supplementary Fig. 8.**
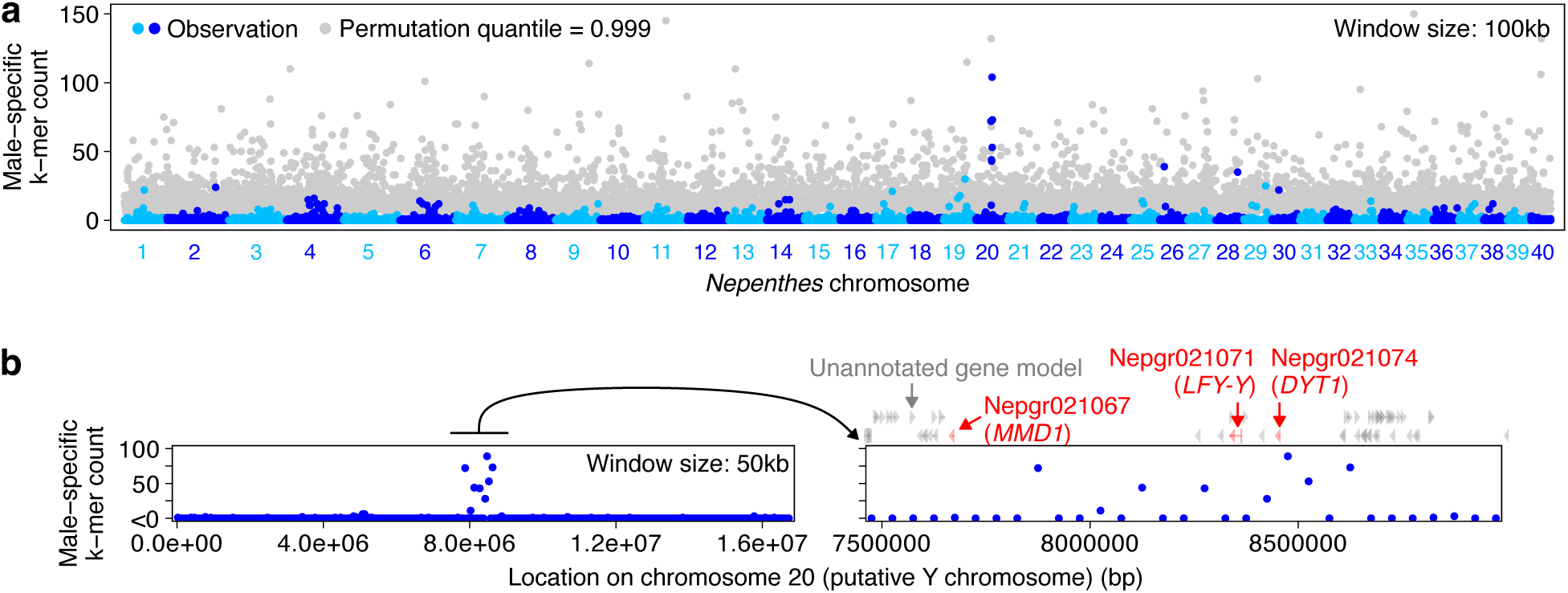
The frequency of male-specific *k*-mers in the *Nepenthes gracilis* male genome. (**a**) The genome-wide distribution of male-specific 16 mers. (**b**) Magnified views of the male-linked region on chromosome 20.

**Supplementary Fig. 9.**
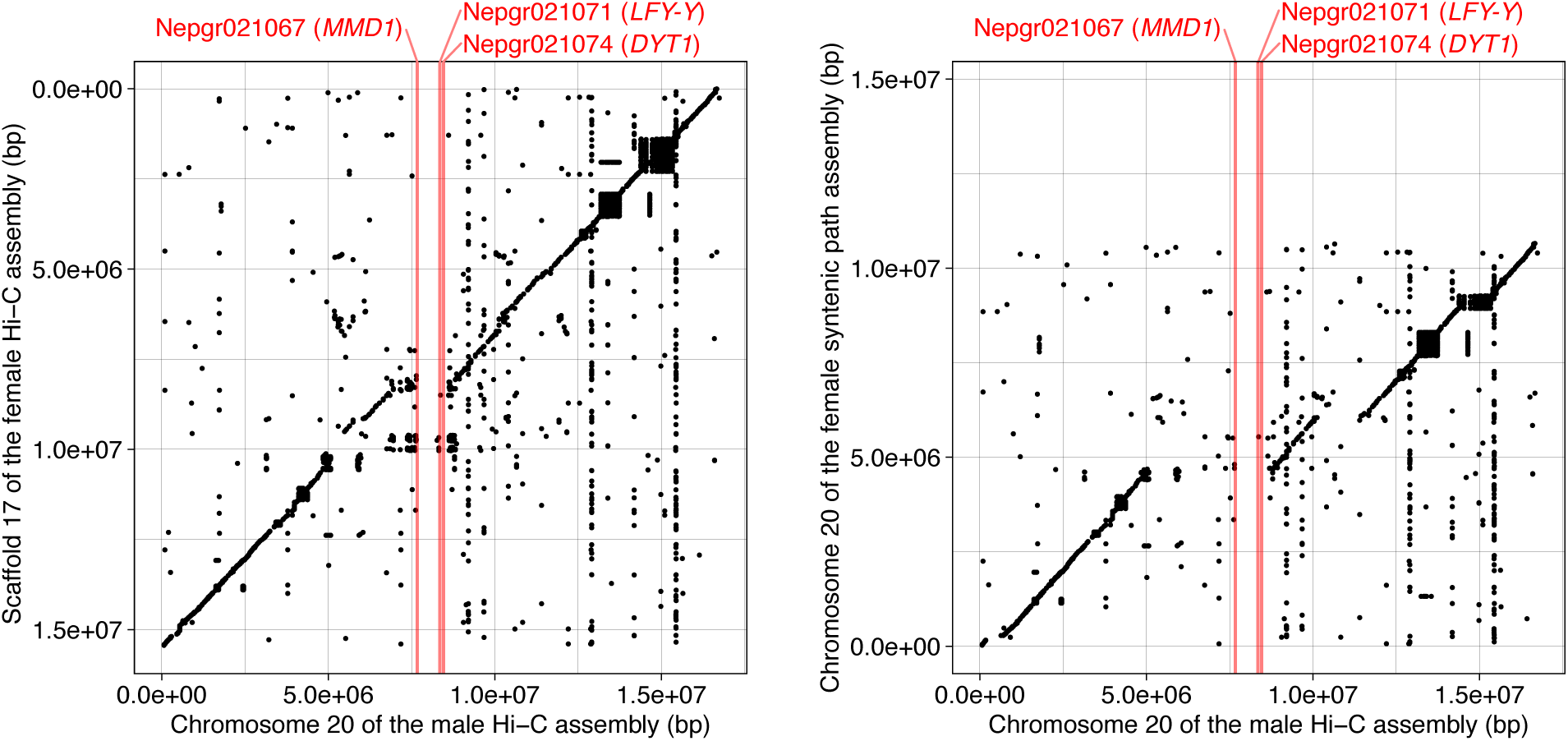
Syntenic comparison of the putative X and Y chromosomes. The female Nanopore assembly was scaffolded with the male Hi-C reads (left) or syntenic path assembly against the male Hi-C assembly (right), and the resulting chromosome-scale female genome assemblies were then used for the dot plot comparison. The male coding sequences were searched against the female genomic sequences using LAST, and all hits are shown in the plot. The positions of the three male-specific transcription factor genes are indicated in red. The results are reproducible using the LAST outputs that are available for download from the following CoGe links: male Hi-C assembly versus female Hi-C assembly (https://genomevolution.org/r/1ir4m) and male Hi-C assembly versus female syntenic path assembly (https://genomevolution.org/r/1nqd6).

**Supplementary Fig. 10.**
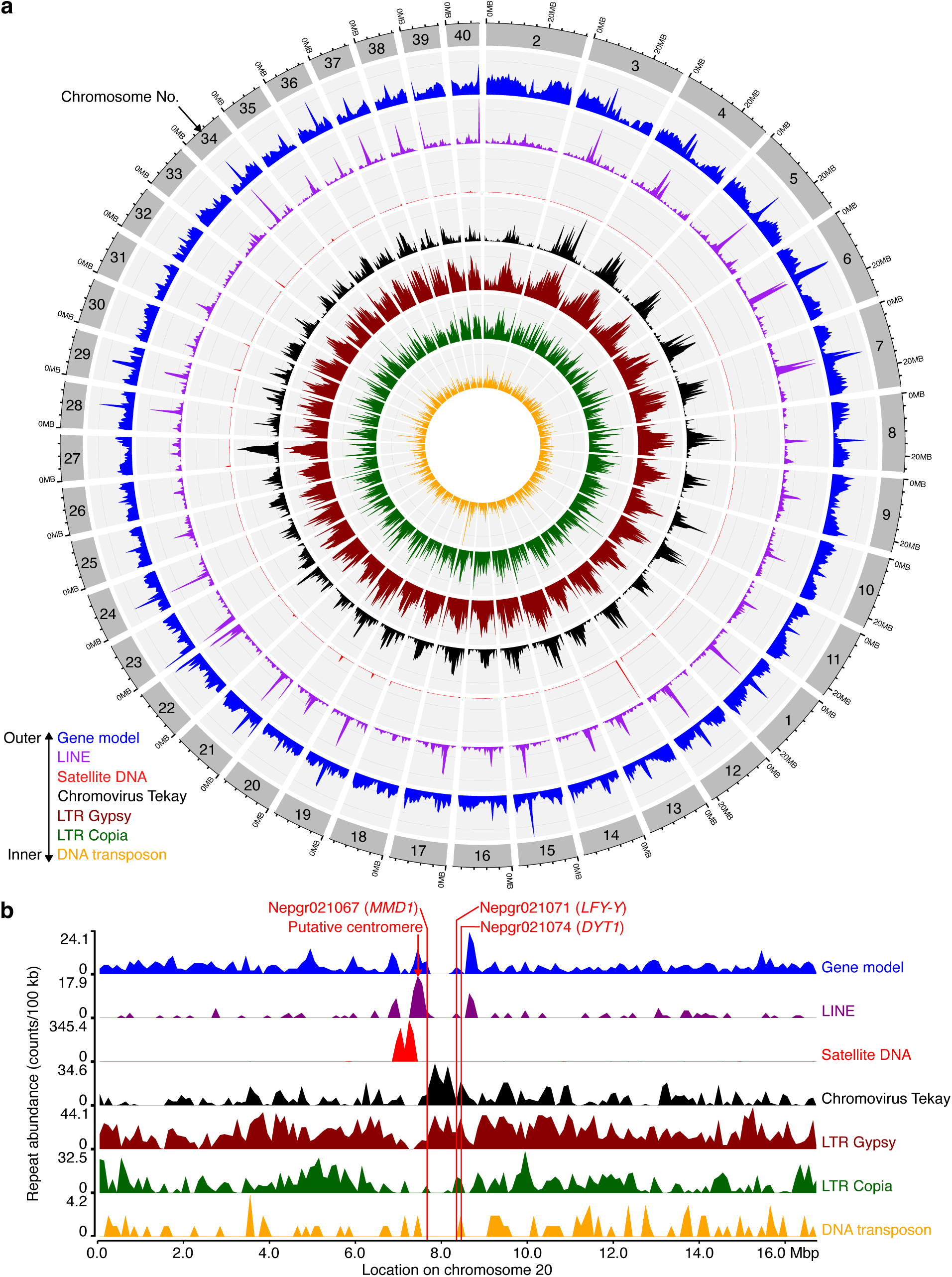
The repeat sequence profiles of the *N. gracilis* male genome. (**a**) Chromosome-wise distributions of representative repeat sequences. Window size: 500 kb. (**b**) A close– up view of chromosome 20. The locations of the putative centromere and male-specific transcription factor genes are indicated. Window size: 100 kb. LINE, long interspersed nuclear elements; LTR, long terminal repeat.

**Supplementary Fig. 11.**
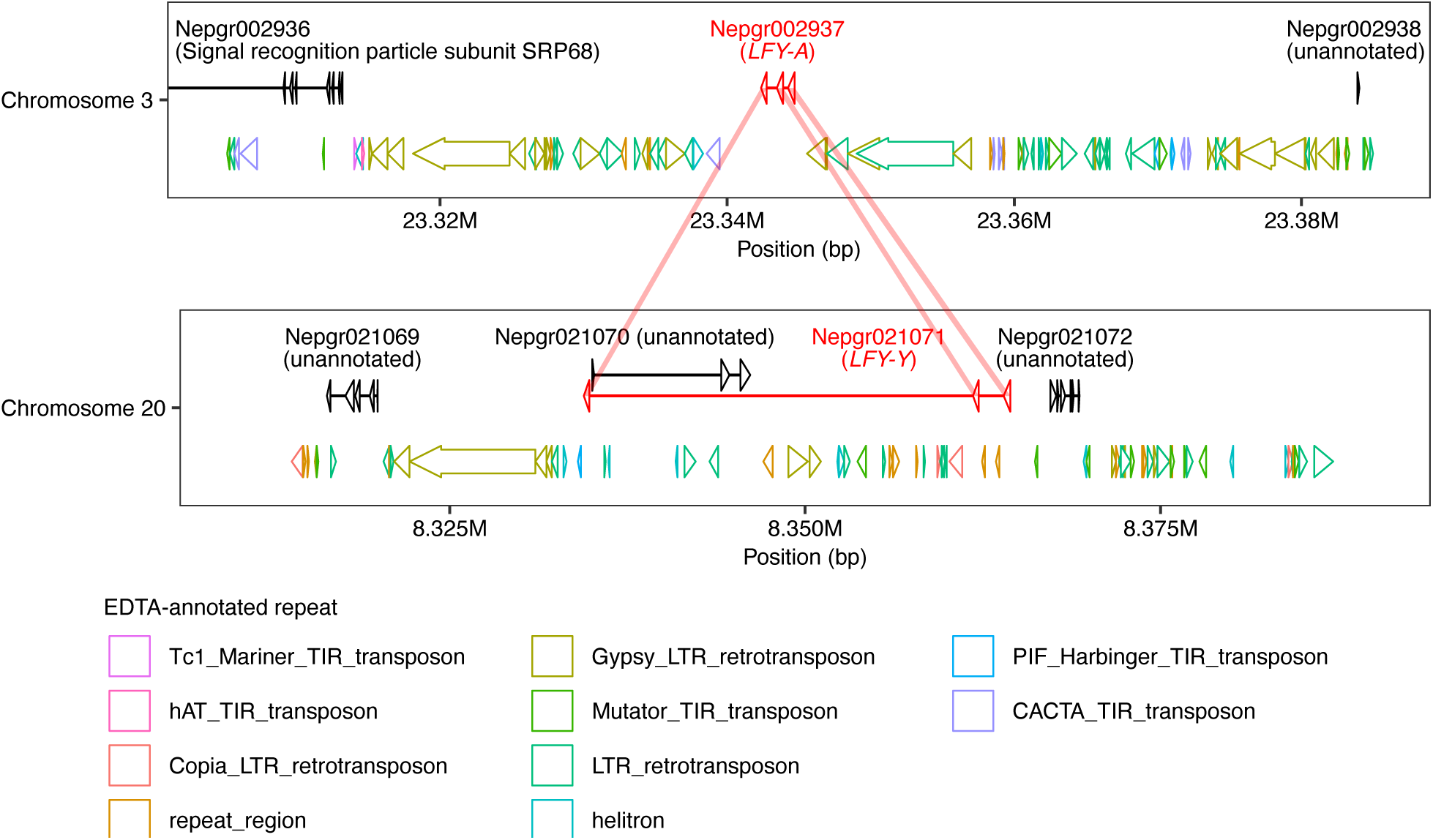
Gene structures of *LFY-A* and *LFY-Y*. Repeat sequences identified by EDTA (Ou et al., 2019) are shown below gene models. Corresponding coding sequences of *LFY-A* and *LFY-Y* are connected. Trinotate-based gene product names are provided in parentheses. The coordinates on chromosomes 3 and 20 have the same scale.

**Supplementary Fig. 12.**
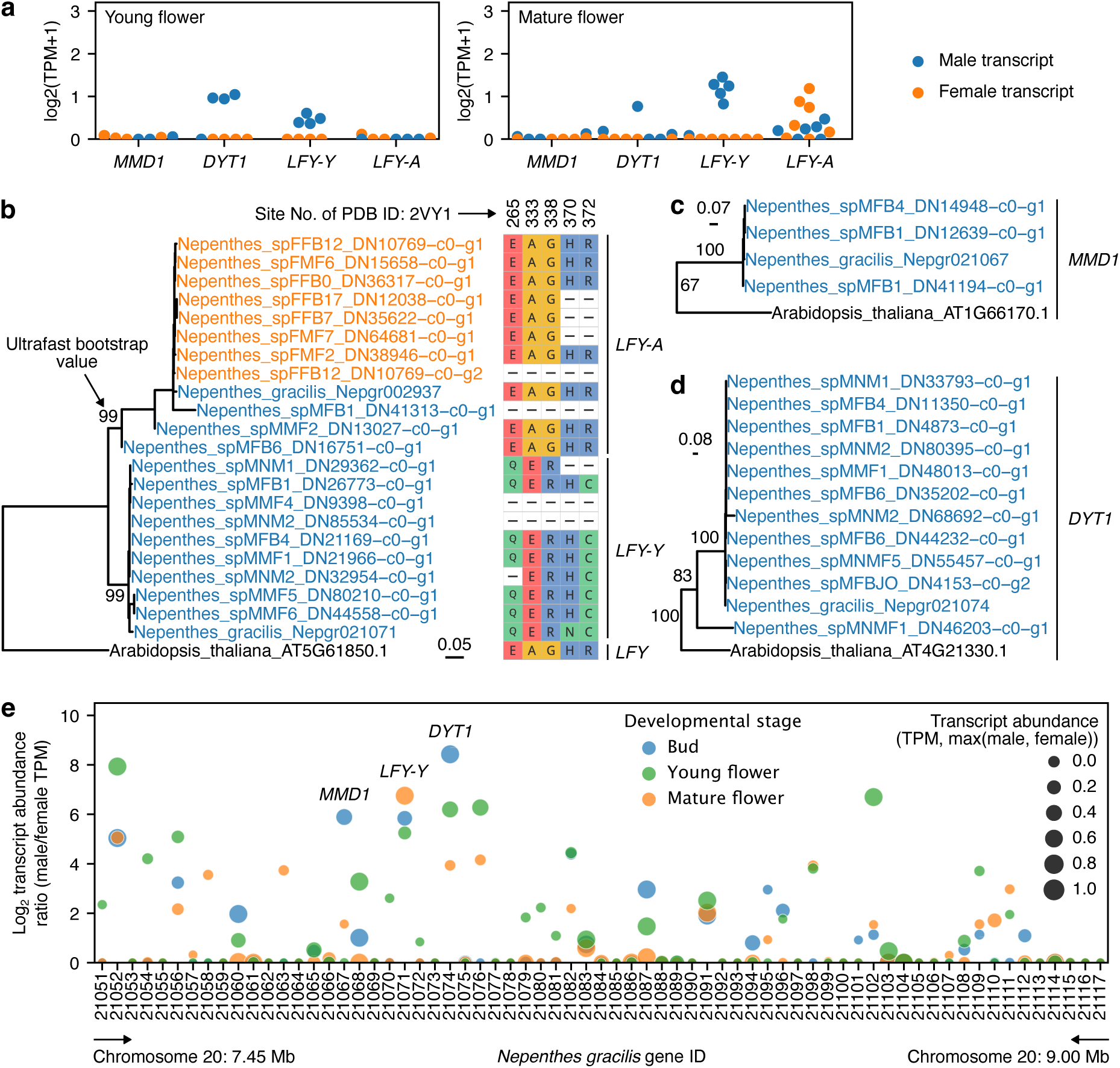
Expression and segregation of male-specific genes. (**a**) The expression levels of male-specific genes in the late developmental stages. RNA-seq reads from flowers in *Nepenthes* cultivars and other species were mapped to the *N. gracilis* male gene models to estimate the transcript abundance as in Fig. 3e. For sample dissection, see Supplementary Fig. 21. (**b**) The phylogenetic identification of *LFY-A* and *LFY-Y* in individually assembled flower transcriptomes. *LFY* transcripts were identified from 17 out of 31 transcriptome assemblies using the TBLASTX search with the *N. gracilis* genes as query sequences with the E-value cutoff of 0.01 and a minimum query sequence coverage of BLAST hits of 25%. The amino acid sites where a substitution occurred in the C_LFY_FLO domain of the *N. gracilis* LFY-Y (Supplementary Fig. 13) are shown to the right. Note that the missing amino acid positions represent the incomplete assembly of the transcript. The clade of *LFY-A* contains transcripts from both male and female individuals (blue and orange, respectively), while that of *LFY-Y* is composed strictly of male transcripts. The bar indicates nucleotide substitutions per nucleotide site. (**c**) The phylogenetic identification of *MMD1* in individually assembled flower transcriptomes. (**d**) The phylogenetic identification of *DYT1* in individually assembled flower transcriptomes. In **c–d**, orthologous clades were extracted from the full trees containing paralogs after midpoint rooting of the ML tree with the GTR+G4 substitution model. (**e**) The transcript abundance ratios of gene models in the YLR.

**Supplementary Fig. 13.**
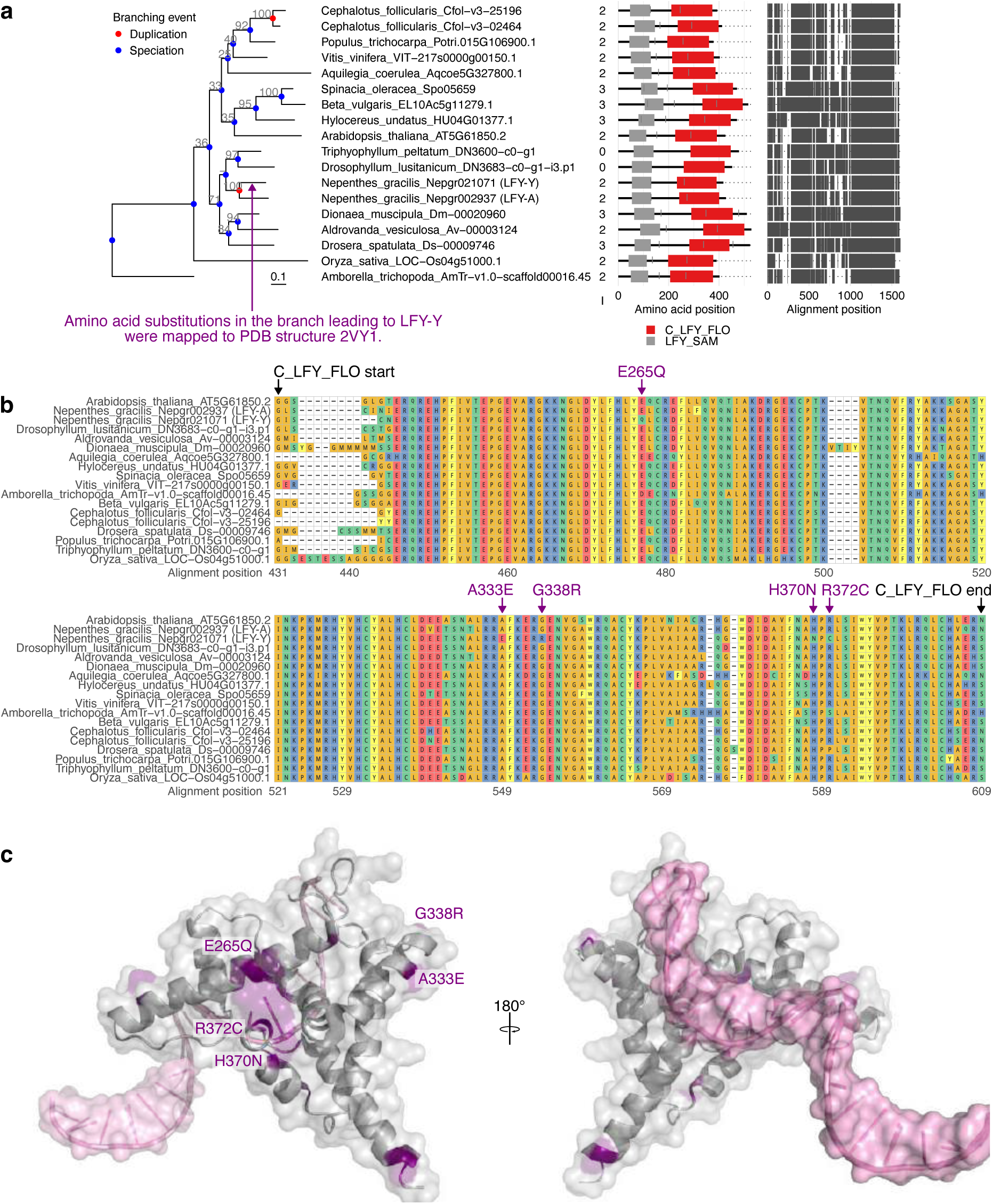
Amino acid substitutions in the conserved C_LFY_FLO domain (pfam17538) of LFY-Y. (**a**) The maximum-likelihood phylogenetic tree of *LFY*. IQ-TREE’s ultrafast bootstrap values are shown above branches. A hyphen (-) marks a branch reconciled by GeneRax. Node colors in the trees indicate inferred branching events of speciation (blue) and gene duplication (red). To the right of the tip labels, the number of introns in protein-coding sequences (I), the Pfam domain structures (E-value < 0.01), and codon alignment structures are shown. Note that the intron numbers are not available in *Ancistrocladus*, *Drosophyllum*, and *Triphyophyllum* as their sequences were obtained from transcriptome assemblies. Both *Nepenthes LFY-A* and *LFY-Y* retain two introns, indicating that the gene duplication was DNA-based. (**b**) The protein alignment of the C_LFY_FLO domain. The positions of amino acid substitutions in the branch leading to LFY-Y are indicated. Note that H370N is not shared by other *Nepenthes* species analyzed in Supplementary Fig. 12, but the others are. (**c**) Amino acid substitutions in *N. gracilis* LFY-Y (purple) mapped to the structure of *A. thaliana* LFY in complex with DNA (pink) from the *APETALA 1* promoter (PDB ID: 2VY1) (Hamès et al., 2008). The substitution mapping was performed using CSUBST (Fukushima and Pollock, 2023).

**Supplementary Fig. 14.**
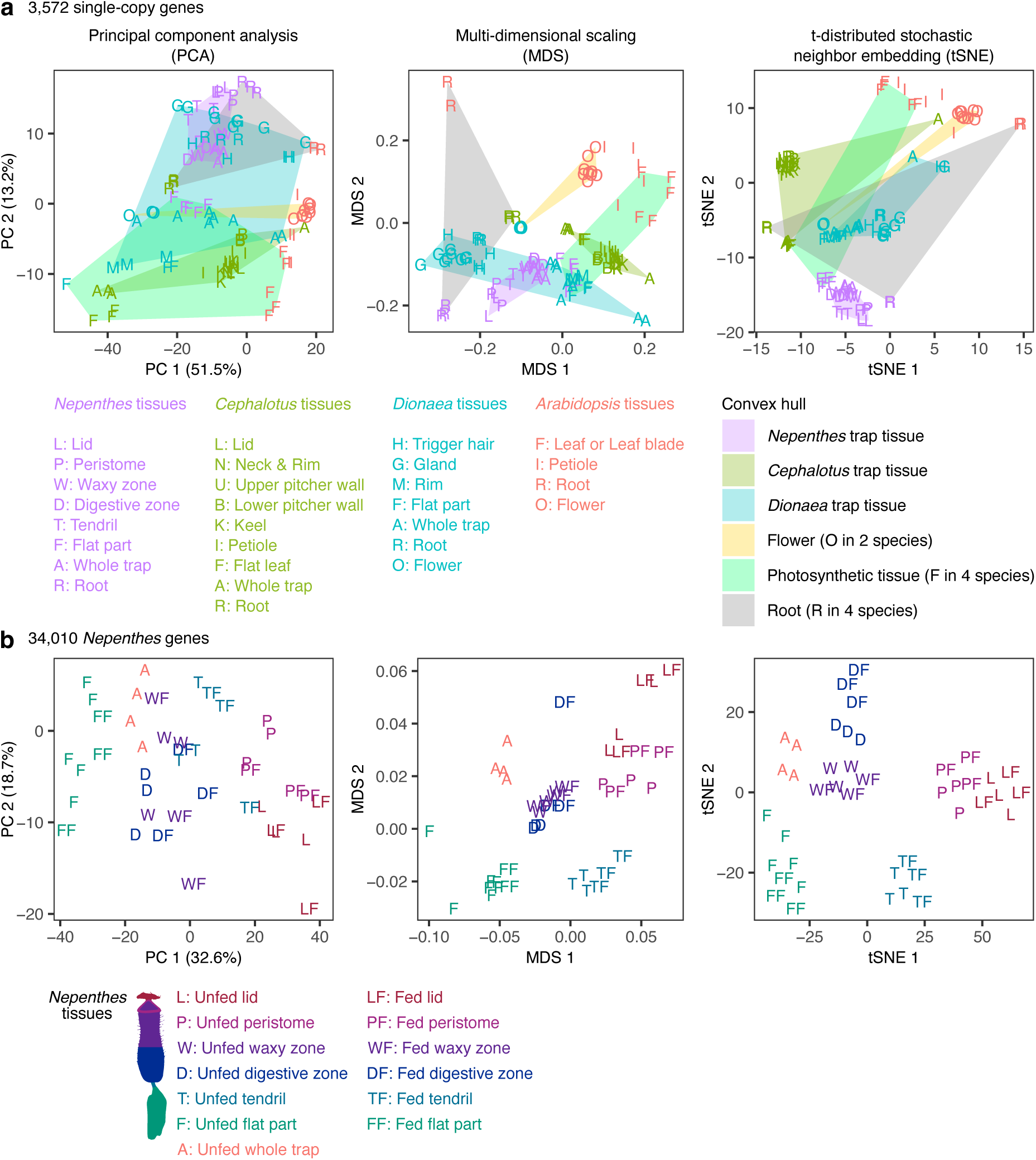
Gene expression profiles of carnivorous trap leaves. (**a**) Comparison among carnivorous trap leaves (*Nepenthes*, *Cephalotus*, and *Dionaea*) and *Arabidopsis* leaves. The cross-species TMM-normalized FPKM values of OrthoFinder-based single-copy genes were analyzed. To draw a robust conclusion, three methods (PCA, MDS, and tSNE) were compared. The distribution of *Nepenthes* trap tissues overlaps with that of roots (PCA) and *Dionaea* trap tissues (PCA, and MDS), but not with others, including the photosynthetic tissues. (**b**) The effect of feeding treatments in the pitcher tissues of *N. gracilis*.

**Supplementary Fig. 15.**
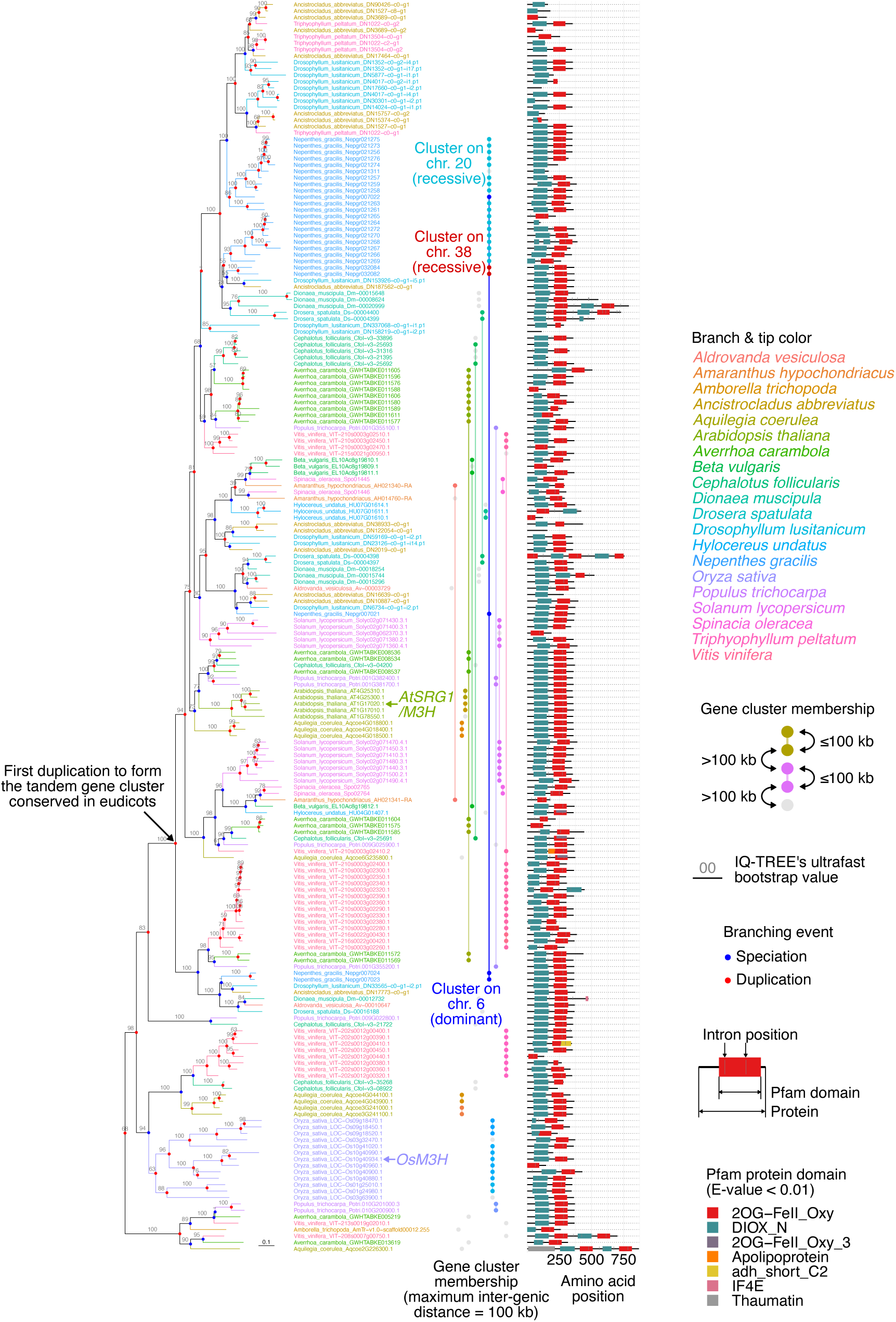
A complete phylogeny of *SRG1*-like genes. Homologous genes were identified by a TBLASTX search with *Arabidopsis* and *Nepenthes* genes as query sequences with the E-value cutoff of 0.01 and a minimum query sequence coverage of BLAST hits of 50%. Phylogenetic analysis was performed as described in Methods. Note that gene cluster memberships could not be obtained for species for which only transcriptomes are available (*Ancistrocladus*, *Drosophyllum*, and *Triphyophyllum*). False negatives of gene cluster detection may occur in species whose genome assemblies are highly fragmented. The bar indicates 0.1 nucleotide substitutions per site.

**Supplementary Fig. 16.**
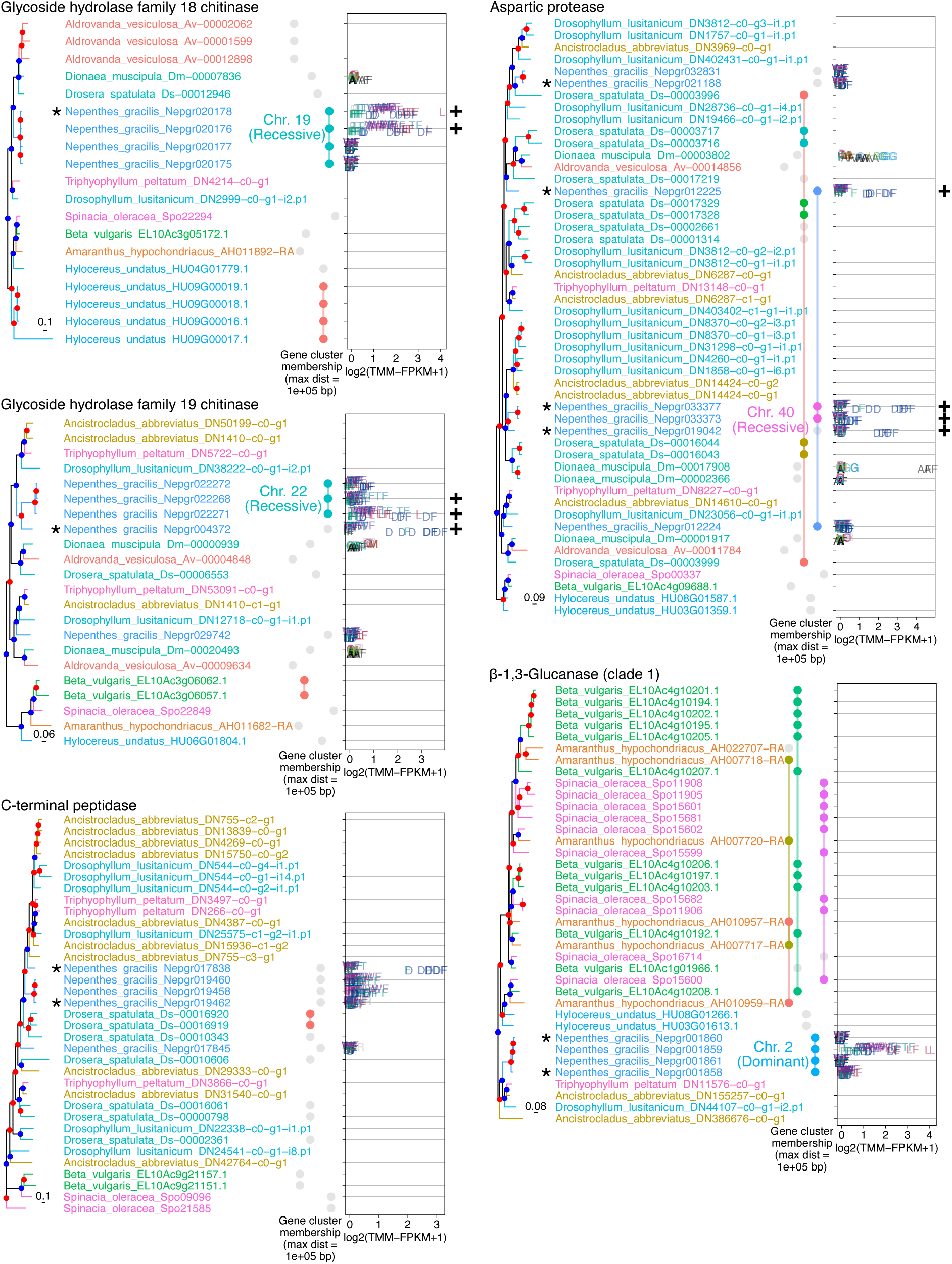
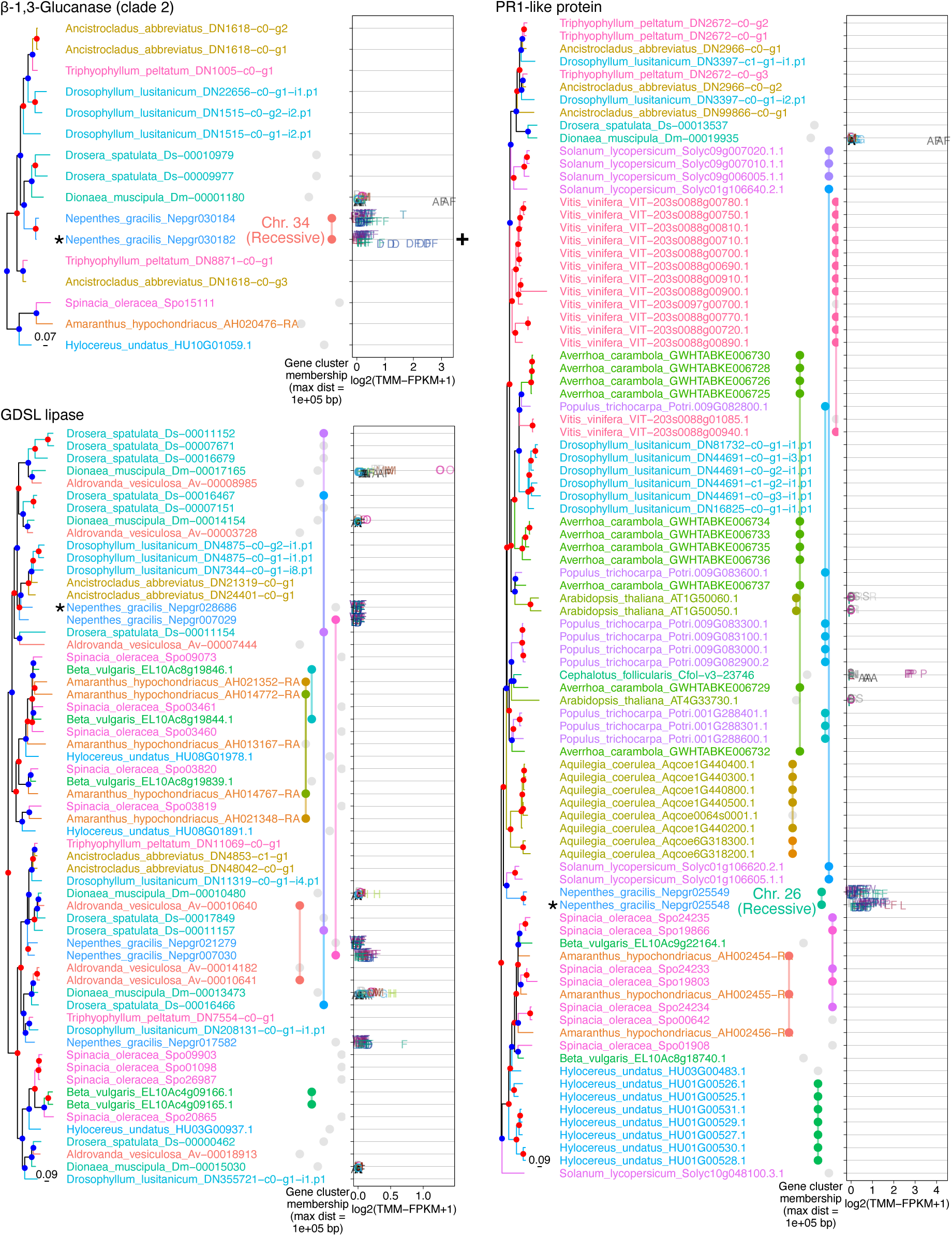
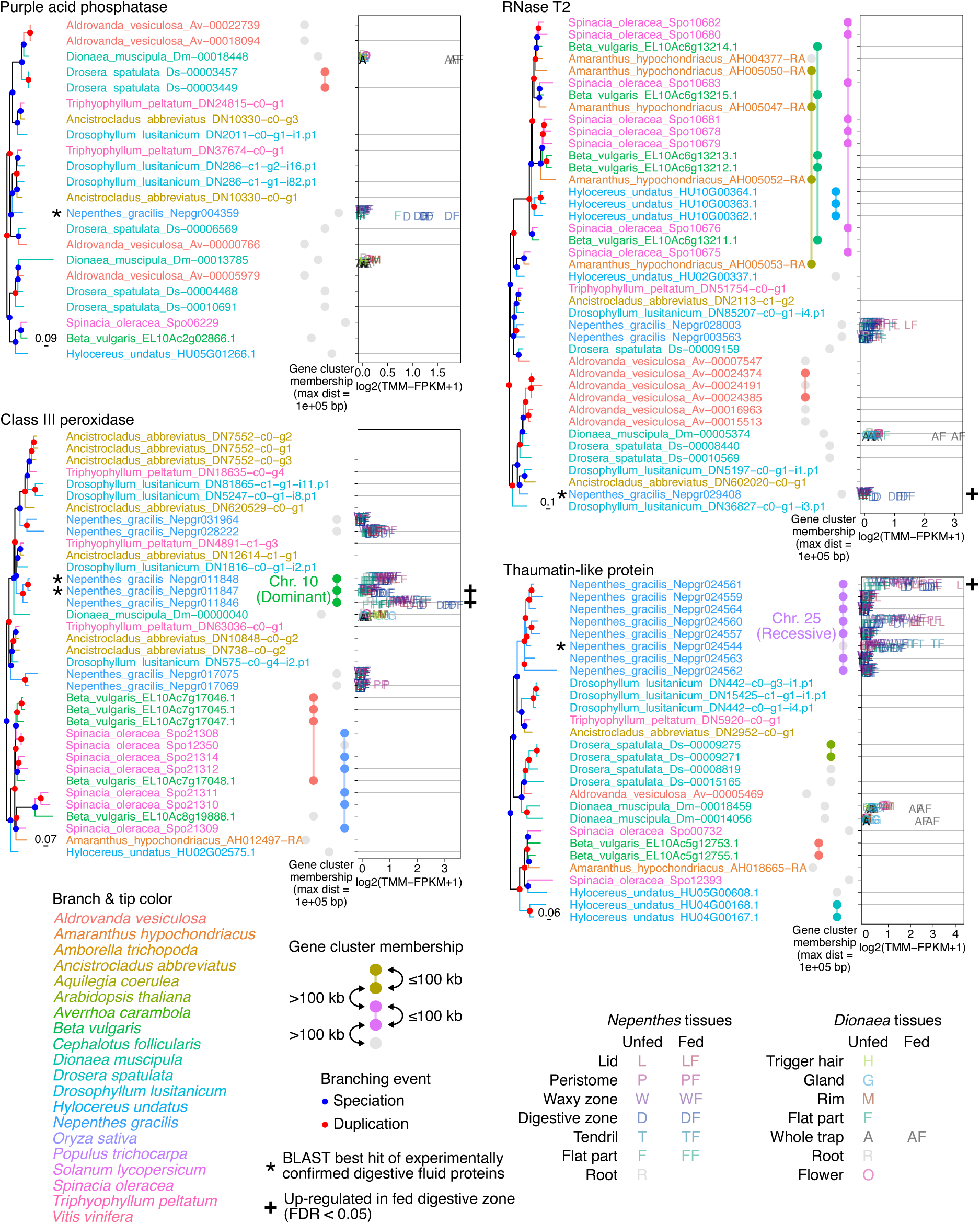
Tandem arrays associated with the genes encoding digestive fluid proteins. The list of experimentally confirmed cases of digestive fluid proteins in *Nepenthes* was obtained from previous work (Fukushima et al., 2017), and their orthologs in the *N. gracilis* genome were identified by a TBLASTX search (asterisks; Supplementary Table 18). Orthologous groups of genes were extracted from the full nucleotide ML trees (Supplementary Dataset). The figure displays, from left to right, the phylogenetic tree, gene IDs, gene cluster memberships, and tissue-specific transcript abundance for each protein family. Transcript abundance is shown for *N. gracilis* and *D. muscipula*. The colors used to represent gene cluster memberships are chosen to differentiate the gene clusters and do not correspond to the colors of the species. Gene clusters associated with digestive fluid proteins are labeled with chromosome IDs and subgenome categories (dominant/recessive).

**Supplementary Fig. 17.**
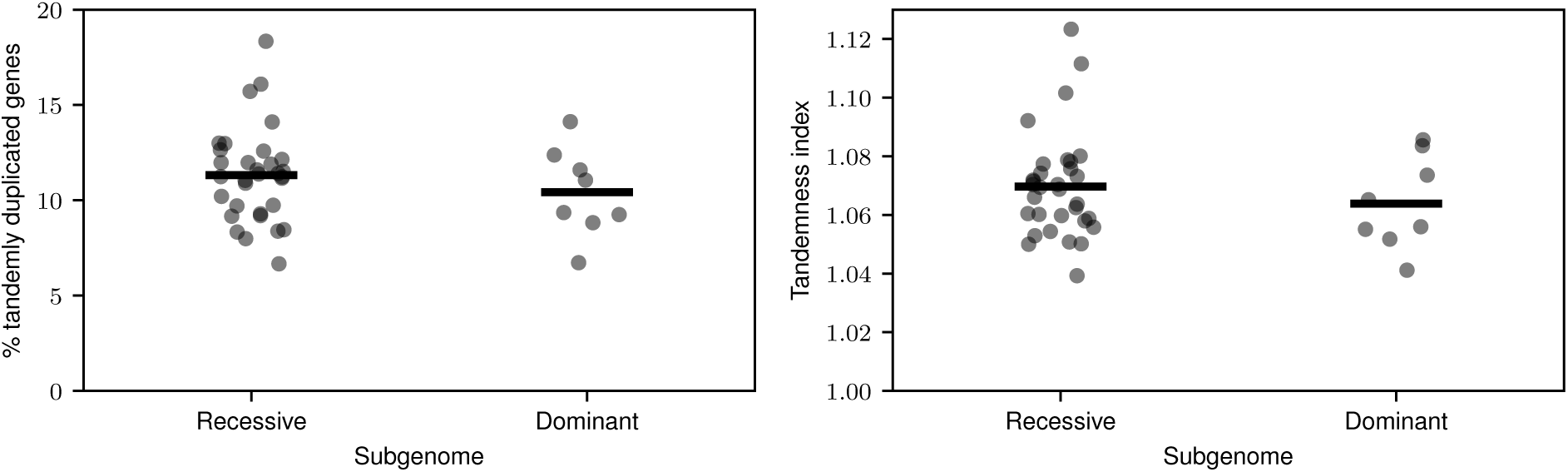
Analysis of chromosome-wise gene tandemness with two different measures. A total of 18,425 *Nepenthes* genes with hits in the UniProt database were analyzed (E-value < 0.01). Tandem duplications were identified as adjacent genes (omitting gene models with no-hit in UniProt) with the same best-hit UniProt entry. The tandemness index was calculated as the mean number of tandem gene groups (including singletons without a tandem duplicate). Points correspond to chromosomes. The numbers of samples are 32 and 8 for recessive and dominant subgenomes, respectively. Bars indicate mean values.

**Supplementary Fig. 18.**
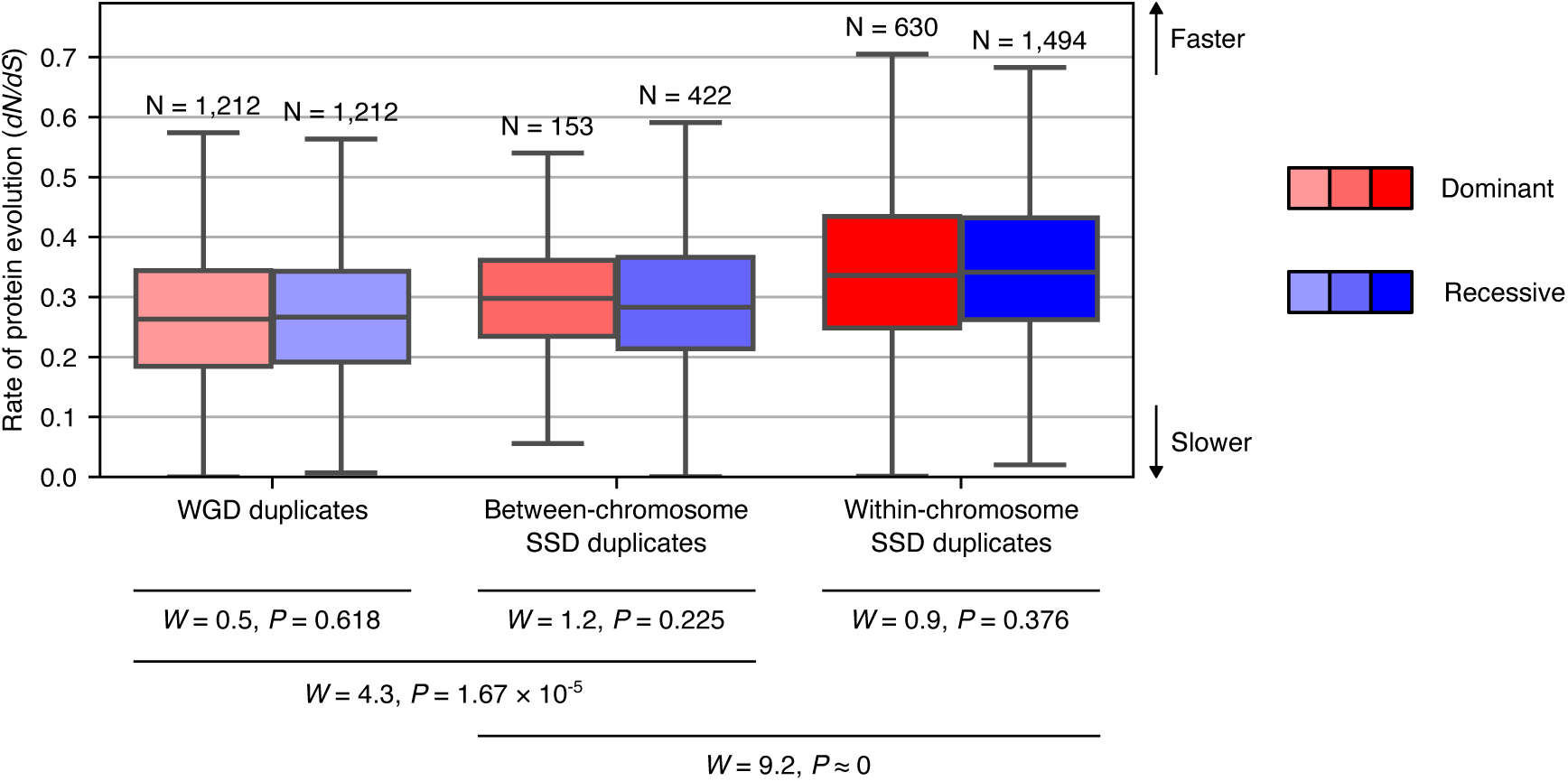
The effect of duplication type and subgenome dominance status on the rate of protein evolution. *dN/dS* was calculated for all branches of reconciled nucleotide ML trees of all orthogroups for the 20 genomes using the *mapdNdS* approach (Guéguen and Duret, 2018), and sister pairs of *N. gracilis* genes were extracted and analyzed. Only expressed genes were included in the analysis (maximum TMM-FPKM > 0.1). Colors match those in Fig. 5. Box plot elements are defined as follows: center line, median; box limits, upper and lower quartiles; whiskers, 1.5 × interquartile range. The stochastic equality of the data was tested by a two-sided Brunner–Munzel test with *W* as the test statistic (Brunner and Munzel, 2000).

**Supplementary Fig. 19.**
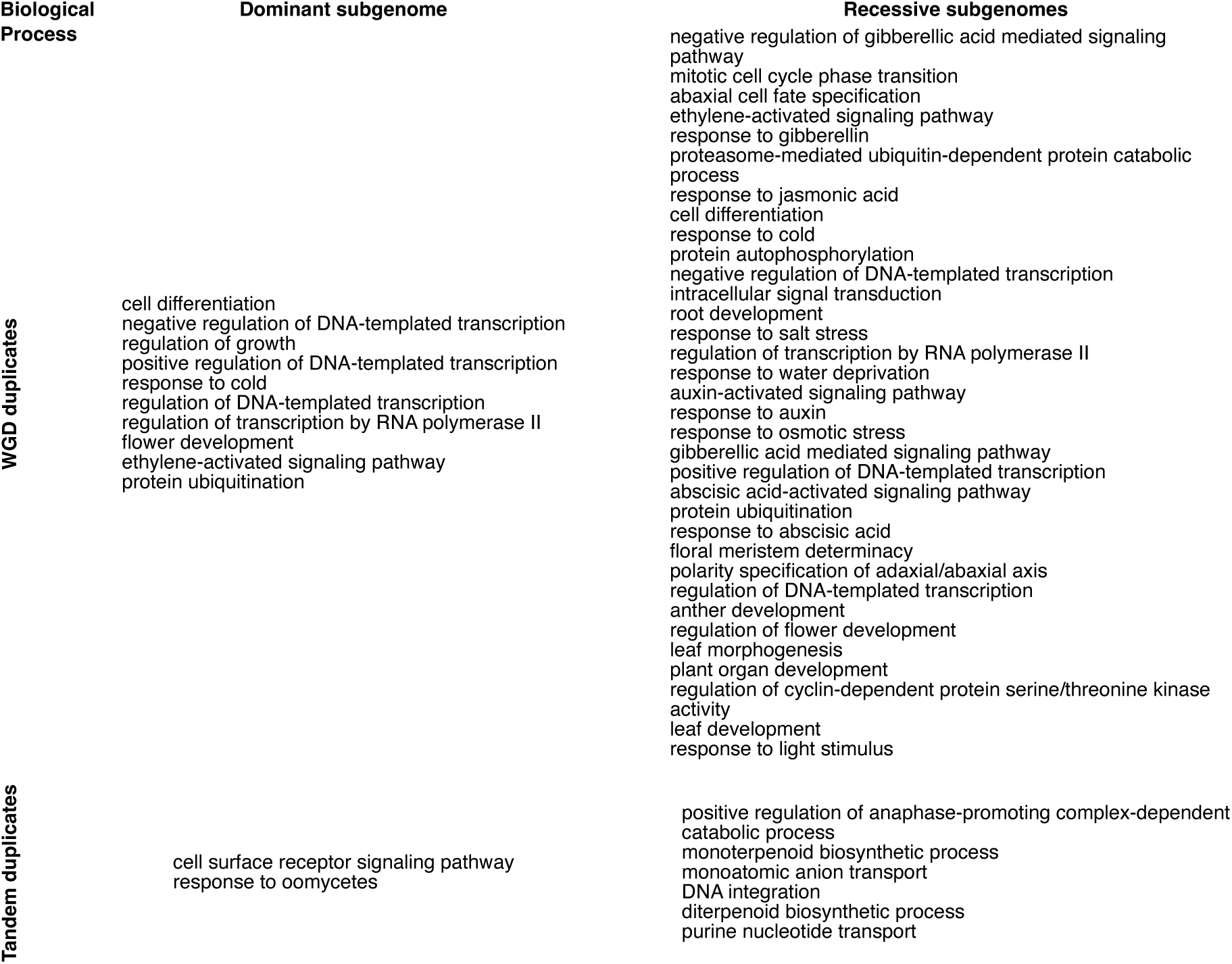
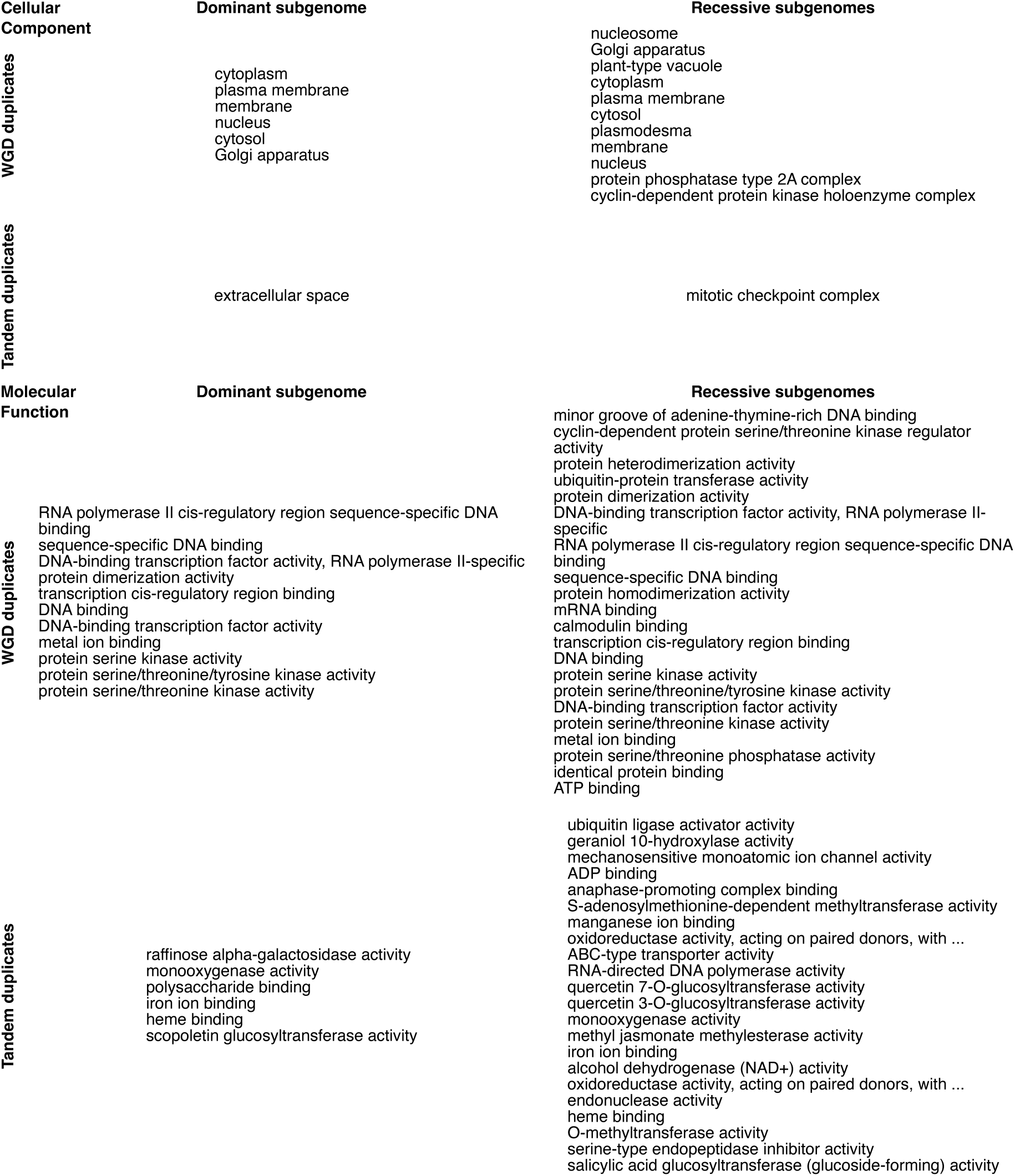
Differential Go enrichments of WGD and tandem duplicates in dominant and recessive subgenomes. For complete information, see Supplementary Table 12, Supplementary Table 13, Supplementary Table 14, and Supplementary Table 15. Following gene annotation, we performed gene ontology (GO) enrichment analysis using GOATOOLS v0.9.9 (Klopfenstein et al., 2018) to test the overrepresentation of gene ontology terms among duplicated genes in dominant chromosomes and recessive chromosomes. The background consisted of all annotated genes in the genome and the foreground subset consisted of syntenic (polyploid) pairs or tandem duplicates, respectively, for both dominant and recessive chromosomes, using Bonferroni-adjusted P < 0.05 as the threshold for significance. Syntenic vs. tandem duplicates were downloaded from a self-vs-self analysis of the *Nepenthes gracilis* genome using CoGe’s SynMap tool (Haug-Baltzell et al., 2017).

**Supplementary Fig. 20.**
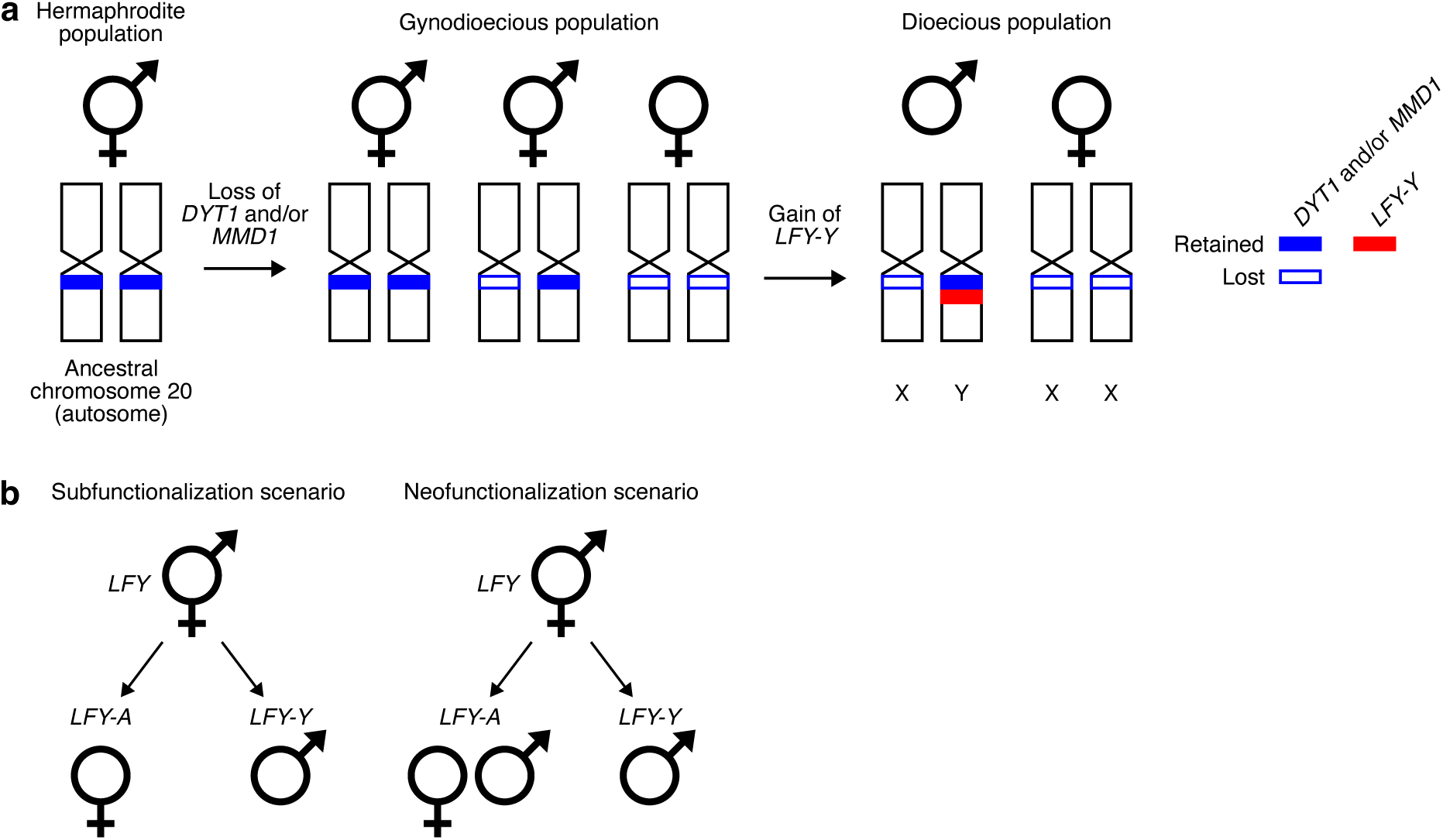
A model for the evolution of dioecy in *Nepenthes*. (**a**) A plausible scenario for the transition from hermaphrodite to dioecious populations. (**b**) Functional divergence after gene duplication of *LFY*.

**Supplementary Fig. 21.**
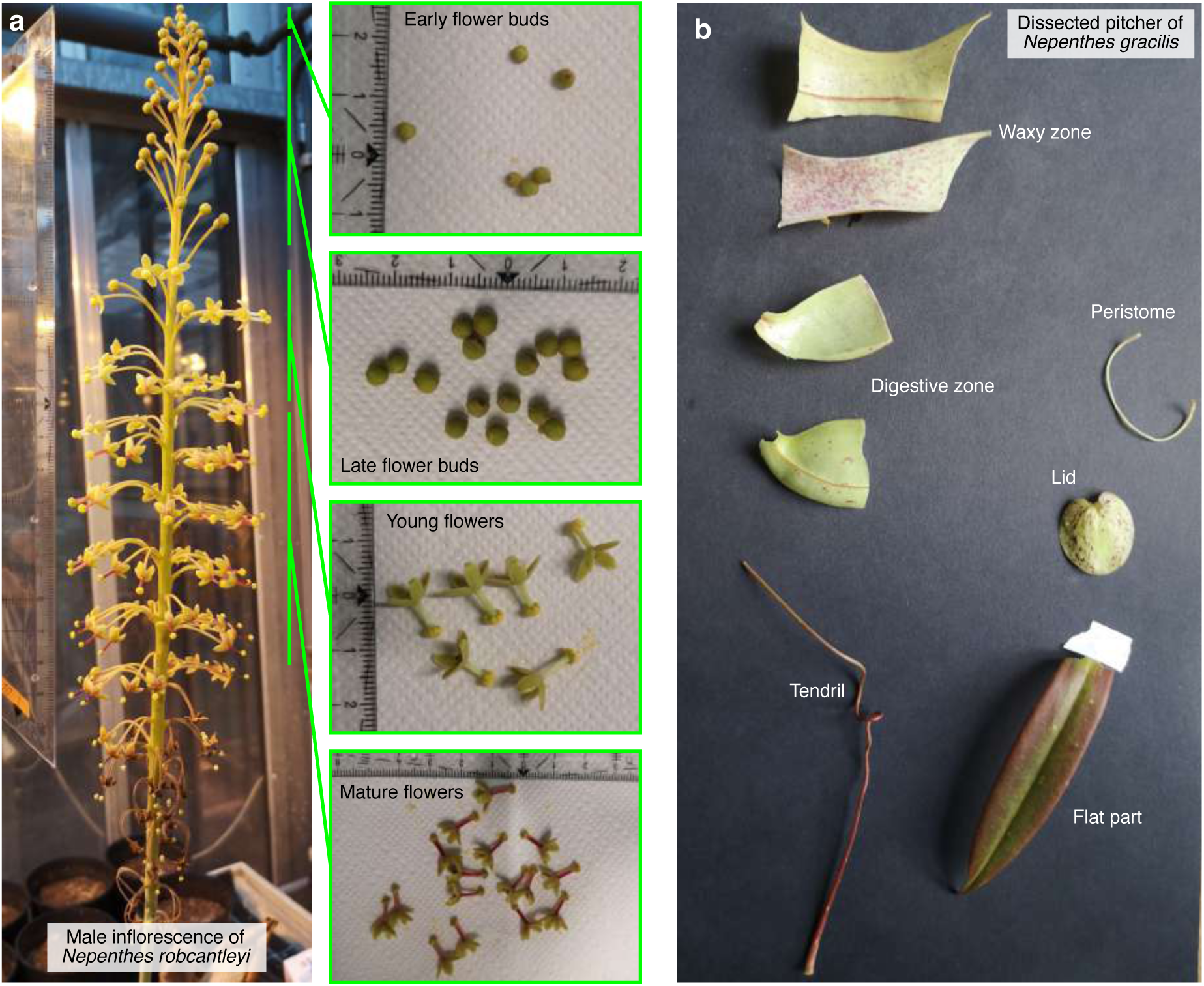
Dissection of *Nepenthes* samples for RNA-seq analysis. (**a**) Inflorescence dissection. The early flower buds included the six youngest flower buds at the tip of the developing inflorescence. In the expression analysis (Fig. 3 and Supplementary Fig. 12), early and late flower buds were not distinguished and were analyzed together as flower buds. Inflorescences were dissected in the same manner regardless of species and sex. (**b**) The pitcher dissection. The transitional part between the digestive zone and the tendril was carefully removed to prevent the cross-contamination of tissues.

**Supplementary Fig. 22.**
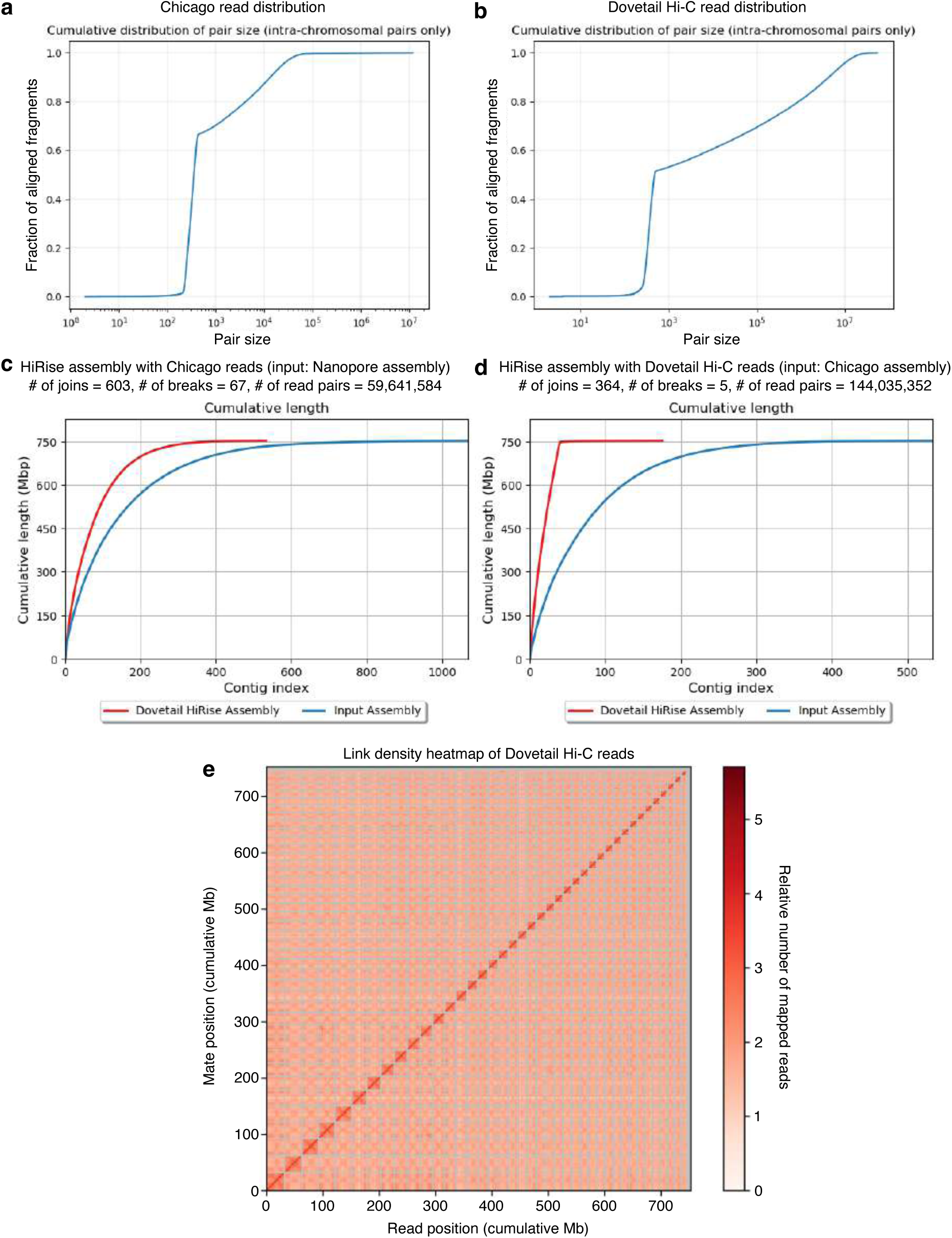
Hi-C assembly of the *N. gracilis* male genome. (**a-b**) The distributions of read pair sizes in Chicago (**a**) and Dovetail Hi-C (**b**) libraries. (**c-d**) Improved contiguity of the genome assembly with Chicago (**c**) and Dovetail Hi-C (**d**) reads. (**e**) The link density heatmap of the Dovetail Hi-C reads. The X-axis and Y-axis represent the mapping positions of the first and second reads in read pairs, respectively. Gray lines indicate the boundaries between scaffolds. The value range of heatmap colors for the number of mapped reads was set to 0–351 in Juicebox (https://github.com/aidenlab/Juicebox).

**Supplementary Fig. 23.**
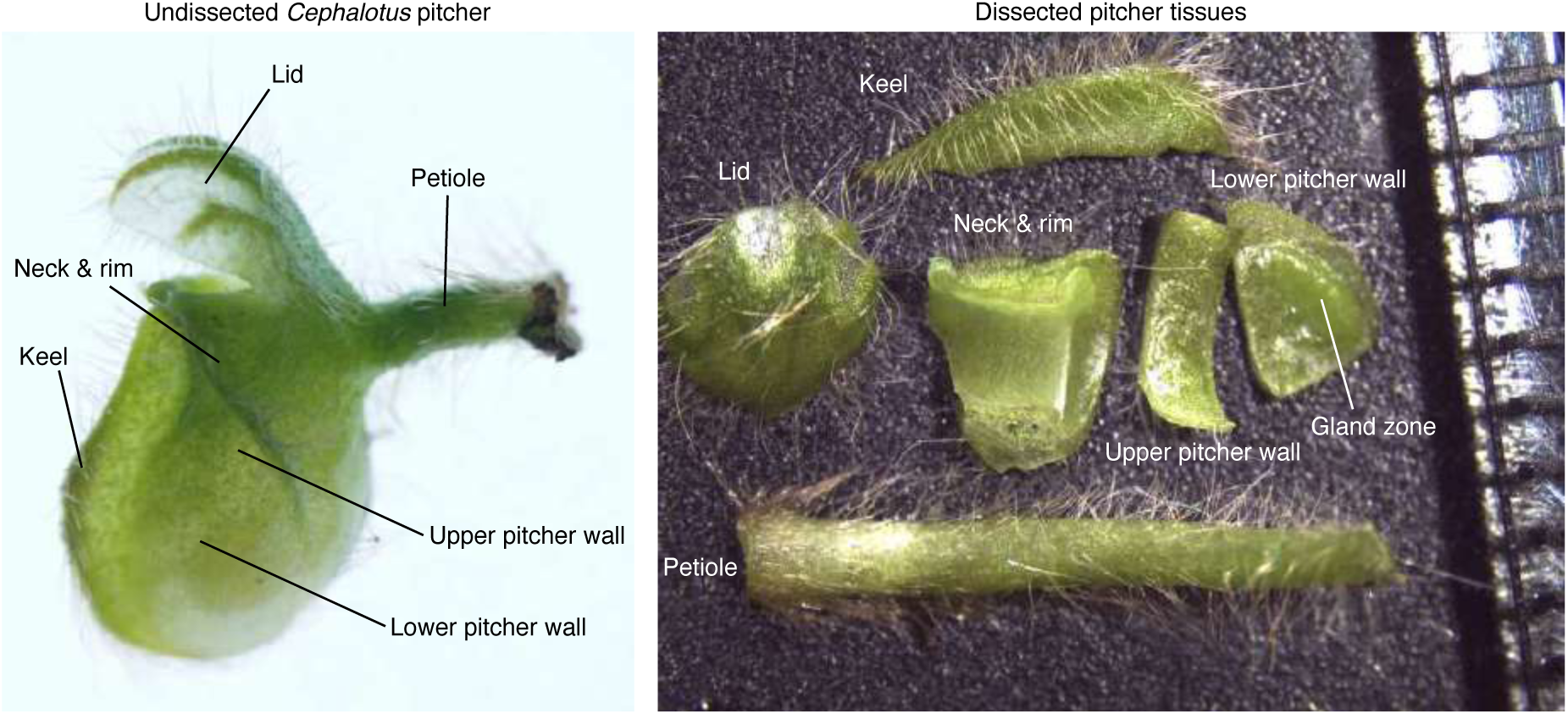
Dissection of pitcher leaves in *Cephalotus follicularis* for RNA-seq analysis. The smallest increment of the ruler is 1 mm.

**Supplementary Fig. 24.**
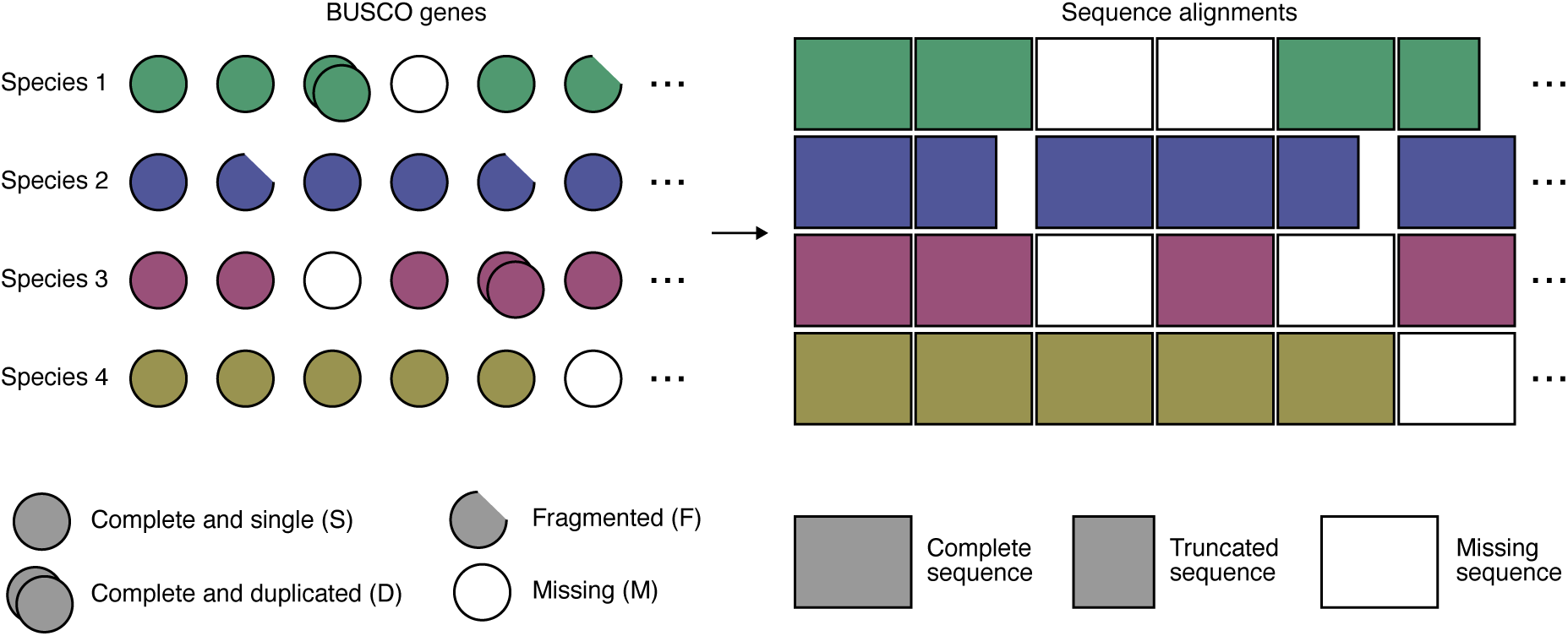
Generating single-copy gene alignments. All detected BUSCO genes marked as single-copy (S) or fragmented (F) were extracted, while those marked as duplicated (D) or missing (M) were treated as missing data.

